# Proteostatic significance of helix O alanine residues in CLC channels

**DOI:** 10.64898/2026.05.25.725697

**Authors:** Chia-Ying You, Ciao-Yu Zhong, Pei-Chen Tsai, An-Ting Cheng, Yu-Xuan Chen, Chung-Jiuan Jeng, Chih-Yung Tang

**Author notes:** Corresponding authors: Dr. Chih-Yung Tang, Department of Physiology, College of Medicine, National Taiwan University, Taipei 100, Taiwan., E-mail address, Dr. Chung-Jiuan Jeng, Institute of Anatomy and Cell Biology, College of Medicine, National Yang Ming Chiao Tung University, Taipei 112, Taiwan.

## Abstract

The voltage-gated chloride channels ClC-1 and ClC-2 are homodimeric structures essential for maintaining muscle excitability and tissue fluid homeostasis, respectively. Mutations in these channels disrupt protein homeostasis (proteostasis), leading to hereditary disorders such as myotonia congenita and leukodystrophy. Specifically, the substitution of highly conserved alanine residues within the transmembrane helix O, exemplified by ClC-1 (A531V) and ClC-2 (A500V), results in severe proteostatic defects characterized by reduced protein stability and impaired surface trafficking. However, the precise role of helix O in these pathological processes remains poorly understood. In this study, we investigated these conserved residues using biochemical and functional approaches. Our findings demonstrate that even subtle structural alterations at these critical sites significantly interfere with channel stability and membrane expression. This study highlights the critical contribution of helix O to proper CLC channel folding and endoplasmic reticulum (ER) quality control, providing deeper insights into the molecular mechanisms of CLC-related channelopathies.

## INTRODUCTION

The voltage-dependent chloride channels ClC-1 and ClC-2 are members of the CLC channel/transporter superfamily. Like other members of the CLC channel/transporter family, functional ClC-1 and ClC-2 channels comprise homodimeric structures, with each monomer containing an intracellular α-helix (helix A), 17 transmembrane α-helices (helices B to R), and two cytosolic cystathionine-β-synthase (CBS) domains, which serve as ATP-binding sites (Jentsch et al., 2002; Tseng et al., 2011; Wang et al., 2019; Xu et al., 2024) (Fig. 1A). The homodimeric architecture of ClC-1 and ClC-2 lead to distinctive “double-barreled” structure, meaning that each subunit can transport Cl⁻ independently. The process by which Cl⁻ passes through each protopore independently of the other subunit is referred to as “protopore gate”. The other gating process, known as the “common gate”, regulates both protopores simultaneously (Chen, 2005; Jentsch & Pusch, 2018; Ma et al., 2023). In humans, ClC-1 is primarily expressed in skeletal muscle and accounts for approximately 70–80% of the resting membrane conductance, and thus plays a crucial role in (Hoxhaj et al., 2012)maintaining the stability of muscle membrane excitability (Bretag, 1987; Dulhunty, 1979). On the other hand, human ClC-2 is widely expressed across nearly all tissues, with particularly high levels in neurons, glial cells, and epithelial cells (Jentsch & Pusch, 2018; Thiemann et al., 1992). The ClC-2 channel contributes to hyperpolarization-induced macroscopic chloride currents with slow activation kinetics (Thiemann et al., 1992).

**Figure 1.**
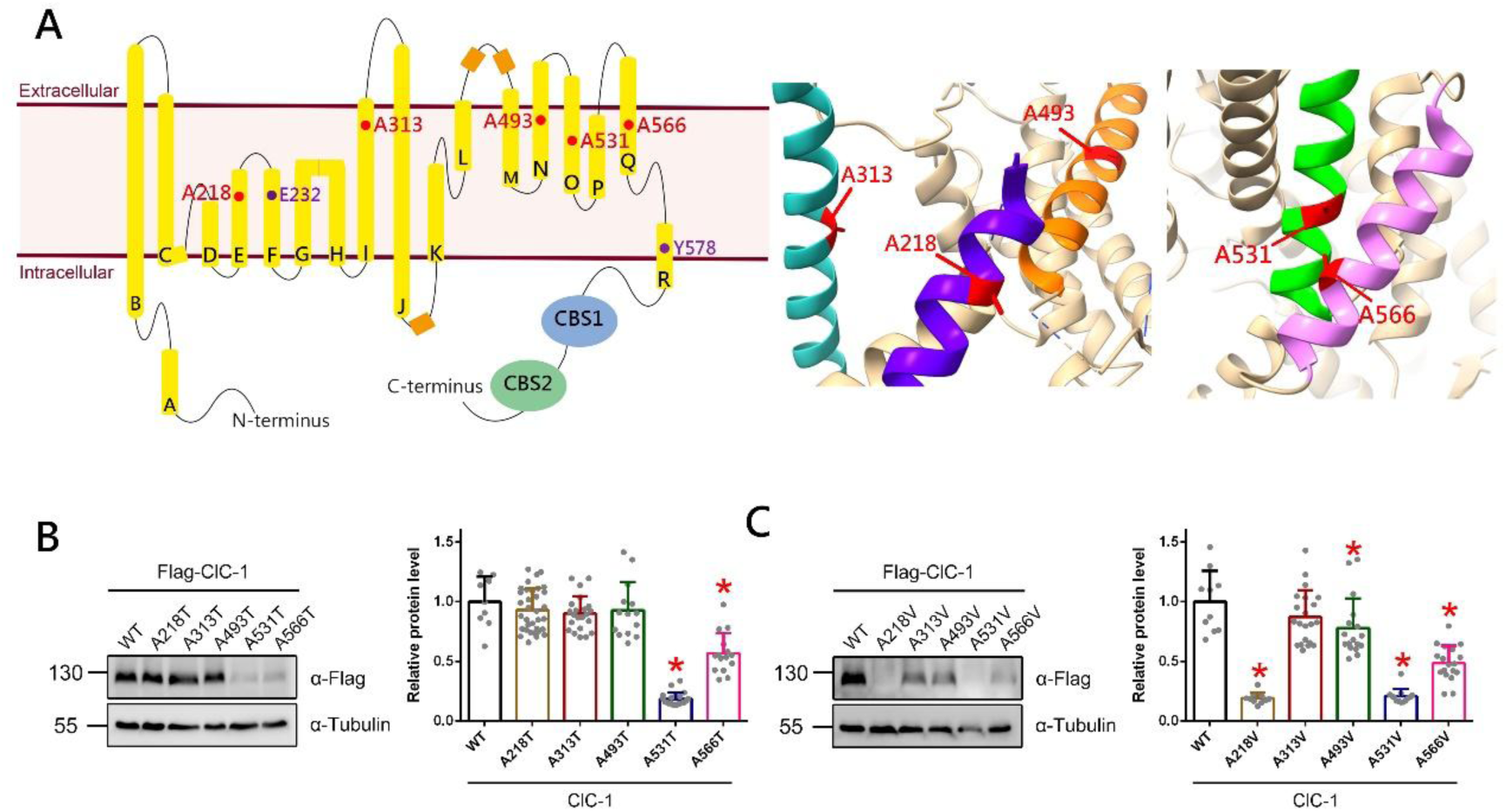
Myotonia congenita-associated ClC-1 alanine mutations. (A) (left) Schematic representation of the membrane topology of the human ClC-1 subunit, with disease-associated mutations highlighted in red. Pore-lining residues are marked in purple. The glutamate gate (E232) regulates each individual protopore by competing with Cl⁻ for binding sites through its negatively charged side chain, thereby controlling Cl⁻ permeation (Dutzler et al., 2003; Park & MacKinnon, 2018; Wang et al., 2019). Tyrosine-578 (Y578), may occupy the Cl⁻ binding site and contribute to the intracellular portion of the selectivity filter (Park & MacKinnon, 2018; Wang et al., 2019). (right)The structure of human ClC-1 subunit (PDB ID: 6COY) was presented using UCSF chimeraX. Highlighted are helices containing myotonia-related residues: helix E (purple), helix I (light blue), helix N (orange), helix O (green), and helix Q (pink). Disease-associated residues are shown in red. (B) Immunoblot analysis and quantification of flag-tagged ClC-1 wild type and alanine to threonine myotonia-associated mutants. Normalized ClC-1 protein level: WT, 1.00±0.21; A218T, 0.93±0.17; A313T, 0.90±0.14; A493T, 0.92±023; A531T, 0.18±0.05; A566T, 0.56±0.16 (*, *P* < 0.05; n = 9–35). (C) Immunoblot analysis and quantification of flag-tagged ClC-1 wild type and alanine to valine myotonia-associated mutants. Normalized ClC-1 protein level: WT, 1.00±0.25; A218T, 0.19±0.04; A313V, 0.87±0.22; A493V, 0.77±0.24; A531V, 0.20±0.06, A566V, 0.48±0.14 (*, *P* < 0.05; n = 10–21).

Mutations in the human gene *CLCN1*, which encodes ClC-1 channels, have been associated with the hereditary skeletal muscle disorder myotonia congenita. Depending on the mode of inheritance, myotonia congenita can be classified as either autosomal dominant (Thomsen’s disease) or autosomal recessive (Becker’s disease) (Ptacek et al., 1993). Specifically, this inherited disease is characterized by delayed muscle relaxation following voluntary contraction. Animal models of myotonia, such as goats and mice, exhibit sustained contractions of the flexor and extensor muscles of the limbs. (Cannon, 2015; Lossin & George, 2008; Pusch, 2002). In addition, the recessive myotonia model *adr* mice displays a marked reduction in skeletal muscle Cl⁻ conductance. Similarly, loss-of-function mutations in *CLCN2* were associated with leukodystrophy, a type of *CLCN2*-related leukoencephalopathy, in which ClC-2 dysfunction leads to fluid accumulation causing myelin vacuolation in mice (Depienne et al., 2013; Wang et al., 2017).

Disease-causing ClC-1 and ClC-2 mutations may manifest aberrant voltage-dependent gating, as well as defective protein homeostasis (proteostasis) (Fu et al., 2020; Gaitan-Penas et al., 2017; Lee et al., 2013; Papponen et al., 2008). Proteostasis refers to the dynamic regulation of protein homeostasis through coordinated mechanisms of protein synthesis and degradation, ensuring proper folding and functional localization within the appropriate subcellular compartments (Balch et al., 2008; Labbadia & Morimoto, 2015). Disease-related ClC-1 and ClC-2 mutations with defective protein stability exhibit reduced protein expression levels or impaired channel trafficking.

A myotonia-causing ClC-1 mutation of an alanine residue in the transmembrane helix O (A531V) has been associated with impaired protein stability (Lee et al., 2013; Papponen et al., 2008). In addition, a mutation of an alanine residue in helix O of ClC-2 (A500V) was linked to leukoencephalopathy and also exhibited significant proteostatic defect (Fu et al., 2021; Gaitan-Penas et al., 2017). Both mutations involve the substitution of highly conserved alanine residues within helix O of ClC channels. Mutations of conserved alanine residues within helix O of ClC channels are implicated in defects of proteostasis, although the underlying mechanisms and the specific role of helix O remain unclear. To explore this issue, we examined these residues through biochemical and functional approaches, revealing that subtle alterations at these sites can interfere with channel stability and surface expression. These findings support the notion that helix O contributes critically to ClC channel folding and quality control in the endoplasmic reticulum.

## MATERIAL and METHODS

### Protein homology modeling

Structural rendering and molecular modeling were carried out with the UCSF ChimeraX (Pettersen et al., 2021). The cryo-EM structures of human ClC-1 (PDB: 6COY) and ClC-2 (PDB: 7XF5) were used as templates for the ClC-1 and ClC-2 channel conformations, respectively (Ma et al., 2023; Park & MacKinnon, 2018). The relative residue positions within the ClC-1 topology were determined by referencing the data established in the previous study (Brenes et al., 2023).

### cDNA constructs

Flag-ClC-1 was generated by subcloning human ClC-1 (NM_000083) into the pFlag-CMV2 vector (Sigma-Aldrich, St. Louis, MO, USA). Human ClC-2 (NM_004366) was subcloned into the pcDNA3-Flag vector (Invitrogen, Carlsbad, CA, USA) to generate the N-terminal Flag-tagged human ClC-2 construct. Mutations in ClC-1 and ClC-2 were introduced by site-directed mutagenesis, followed by verification with DNA sequencing. Other cDNA constructs employed in this study include pcDNA3-HA rat cereblon (kindly provided by Dr. Chul-Seung Park, Gwangju Institute of Science and Technology, Korea), pCMV3-myc human RNF5 (Sino Biological, Beijing, China).

### Cell culture and DNA transfection

Human embryonic kidney (HEK) 293T cells were grown in Dulbecco’s modified Eagle’s medium (DMEM) (Gibco, Gaithersburg, MD, USA) supplemented with 10% heat-inactivated fetal bovine serum (Hyclone, Logan, UT, USA), 2 mM glutamine, 100 units/mL penicillin and 50 μg/mL streptomycin (Thermo Fisher Scientific, Waltham, MA, USA), and were maintained at 37。C in a humidified incubator with 95% air and 5% CO2. Transient transfection was performed by using the Lipofectamine 2000 reagent (Invitrogen) or T-Pro NTR II transfection reagent (T-pro biotechnology, New Taipei City, Taiwan). Cells were plated in 6-well or 12-well plates 24 h prior to transfection. For specific experiments, cells were seeded onto poly-D-lysine (PDL; Sigma-Aldrich)-coated coverslips in 24-well plates 24 h before transfection. Various expression constructs were incubated with the transfection reagent for 20 min at room temperature, after which DNA–Lipofectamine or DNA–T-Pro NTRII complexes, diluted in Opti-MEM (Gibco) or serum-free DMEM, were added to the culture wells. Cells were harvested 24-48 hrs after transfection. Where indicated, transfected HEK293T cells were incubated in the presence of cycloheximide (Sigma-Aldrich; 100 μg/ml), MG132 (Sigma-Aldrich; 10 μM).

### Co-Immunoprecipitation and Immunoblotting

Transfected cells were incubated at 37°C in the presence of 10 μM MG132 for 24 h. Cells were solubilized in ice-cold immunoprecipitation (IP) buffer [(in mM) 100 NaCl, 4 KCl, 2.5 EDTA, 20 NaHCO3, 20 Tris-HCl, pH 7.5, 1 DTT, 1 PMSF, 1% Triton X-100] containing the protease inhibitor cocktail (Roche Applied Science, Penzberg, Germany). Insolubilized materials were removed by centrifugation. Solubilized lysates were pre-cleared with protein G sepharose beads (GE Healthcare Biosciences, Piscataway, NJ, USA) for 1 h at 4°C, and then incubated for 16 h at 4 °C with protein G sepharose beads pre-coated with the anti-Flag, anti-HA or anti-Myc antibody. Beads were gently spun down and washed twice in a wash buffer [(in mM) 100 NaCl, 4 KCl, 2.5 EDTA, 20 NaHCO_3_, 20 Tris-HCl, pH 7.5] supplemented with 0.1% Triton X-100, and then twice with the wash buffer. The immune complexes were eluted from the beads by heating at 70°C for 5 min in the Laemmli sample buffer.

Protein samples were separated by 7.5–12% SDS-PAGE, electrophoretically transferred to nitrocellulose membranes, and detected using rabbit anti-Flag (1:5000; Sigma-Aldrich), rabbit anti-glyceraldehyde-3-phosphate dehydrogenase (GAPDH) (1:5000; GeneTex, Irvine, CA, USA), rat anti-HA (1:5000; Roche Applied Science), mouse anti-Myc (1:5000; clone 9E10), or mouse anti-tubulin (1:5000; Abcam, Cambridge, UK) antibodies. Blots were then exposed to horseradish peroxidase-conjugated anti-mouse/rabbit IgG (1:5000; Jackson ImmunoResearch,West Grove, PA, USA), and revealed by an enhanced chemiluminescence detection system (Revvity, Waltham, MA, USA). Acquisition of chemiluminescent signals from immunoblots was achieved by using the UVP AutoChemi image system (Ultra-Violet Products, Upland, CA, USA).

For quantitative analysis, data were obtained from at least three independent experiments performed in duplicate or triplicate. Densitometric analysis of immunoblots was carried out using ImageJ (National Institutes of Health, Bethesda, MD, USA). For each blot containing multiple lanes corresponding to the same experimental condition, the signal intensity of the protein of interest was first normalized to that of the corresponding loading control. Normalized values from all control groups were then averaged to obtain the mean control protein density. Each lane, including both treatment and control groups, was subsequently normalized to this mean control value. For conditions tested in multiple experiments, normalized protein density values from individual blots were pooled for statistical analysis.

### Cycloheximide (CHX) chase

To assess protein stability, de novo protein synthesis was blocked in HEK293T cells 24 h post-transfection using cycloheximide (Sigma-Aldrich; 100 μg/ml). Cells were subsequently harvested at the indicated time points (0–6 h or 0–9 h) for immunoblotting analysis. Degradation rates of CLC mutants were determined by standardizing CLC protein density to tubulin signals across the CHX treatment period. All values were subsequently normalized to the 0-h time point to facilitate comparison of protein stability among different variants. To quantify degradation kinetics, a linearized semi-logarithmic plot was used to calculate the half-life of CLC variants. Individual half-life estimates from multiple replicates were then subjected to statistical testing to determine significant differences between experimental conditions.

### Cell surface biotinylation

Cells were washed with ice-cold D-PBS [(in mM) 136 NaCl, 2.5 KCl, 1.5 KH_2_PO_4_, 6.5 Na_2_HPO_4_, pH 7.4, 0.9 mM CaCl_2_ and 0.5 mM MgCl_2_] supplemented with 0.5 mM CaCl₂ and 2 mM MgCl₂, and then incubated with 1 mg/mL sulfo-NHS-LC-biotin (Thermo Fisher Scientific) in D-PBS at 4 °C for 1 h with gentle agitation on an orbital shaker. The biotinylation reaction was quenched by removing the reagent, followed by one wash with 100 mM glycine in PBS and three additional washes with Tris-buffered saline [(in mM) 20 Tris-HCl, 150 NaCl, pH 7.4]. Cells were lysed in lysis buffer [(in mM) 150 NaCl, 50 Tris-HCl, 5 EDTA, 1 PMSF, 1% Triton X-100, pH 7.6] supplemented with a protease inhibitor cocktail. Insoluble material was removed by centrifugation at 4 °C, and the supernatants were incubated overnight at 4 °C with streptavidin-agarose beads (Thermo Fisher Scientific). The beads were washed once with lysis buffer, twice with a high-salt buffer [(in mM) 500 NaCl, 50 Tris-HCl, 5 EDTA, 0.1% Triton X-100, pH 7.6], and once with a low-salt buffer [(in mM) 10 Tris-HCl, 2 EDTA, pH 7.6, 0.1% Triton X-100]. Biotin–streptavidin complexes were eluted by heating the beads at 70°C for 5 min in Laemmli sample buffer. For SDS-PAGE, the amount of lysate loaded in the *Total lane* corresponded to approximately 8% of that used for the streptavidin pull-down and subsequently loaded in the *Surface lane*. To determine surface protein density, surface signals were normalized against total protein levels and subsequently expressed relative to the control group.

### Electrophysiological recordings

Conventional cell-attached and whole-cell patch-clamp techniques were used to record ClC-1 Cl⁻ currents in transfected HEK293T cells. For cell-attached recordings, cells were bathed in an external solution [(in mM) 130 KCl, 5 MgCl₂·6H₂O, 1 EGTA, 10 HEPES, pH 7.4]. Recording pipettes were filled with an internal solution [(in mM) 140 NaCl, 4 CsCl, 2 MgCl₂·6H₂O, 2 CaCl₂, 10 HEPES, pH 7.4]. For whole-cell recordings, the bath solution contained [(in mM) 140 NaCl, 4 CsCl, 2 MgCl₂, 2 CaCl₂, 10 HEPES, pH 7.4], and the pipettes were filled with an internal solution [(in mM) 120 CsCl, 10 EGTA, 10 HEPES, pH 7.4]. Data were acquired and digitized with Axopatch 200A and Digidata 1322A, respectively, via pCLAMP 10 (Molecular Devices, San Jose, CA, USA). Cell capacitances were measured using a built-in function of pCLAMP 10 and were compensated electronically with Axopatch 200A. The holding potential was set at 0 mV. Data were sampled at 10 kHz and filtered at 1 kHz. All recordings were performed at room temperature (20–22°C).

Electrophysiological analyses were performed to determine the voltage dependence of the channel’s open probability (P_o_-V curve). To quantify the open probability (P_o_), the initial tail current amplitude was first determined by fitting the tail current decay with a double-exponential function. This initial tail current value was then normalized to the maximum initial tail current elicited at the most depolarized test potential. Data points in the P_o_-V curve were fitted with a Boltzmann equation: P_o_ =P_min_+(1-P_min_)/{1+exp[zF(V-V_0.5_)/k]} where V_0.5_ is the half-activating voltage for the P_o_-V curve and k denotes the slope factor. We substituted V_0.5_ and k into two formulas: ΔG_0_ mV = zF V_0.5_ and z = (RT/F) k^−1^, calculating the free energy (ΔG_0_ mV) of channel at 0 mV.

Conventional two-electrode voltage clamp (TEVC) experiments were conducted in Xenopus laevis oocytes to record ClC-2 Cl- current. For Xenopus oocyte expression, Flag-hClC-2 cDNA was inserted into the pGEMHE vector (Liman et al., 1992). Subsequently, cDNA templates were linearized and used for in vitro transcription of capped cRNAs using the mMessage mMachine T7 kit (Thermo Fisher Scientific). Xenopus laevis oocytes were obtained from ovarian follicles following an animal protocol approved by the Institutional Animal Care and Use Committee (IACUC) of National Taiwan University. After several washes in collagenase-free, Ca^2+^-free ND96 solution, oocytes were maintained in standard ND96 [(in mM) 96 NaCl, 2 KCl, 1.8 MgCl2, 1.8 CaCl2, 5 HEPES, pH 7.2]. Stage V-VI oocytes were microinjected with human ClC-2 cRNAs and stored at 16 °C for 2-3 days. Prior to recording, injected oocytes were preincubated with 1 percent dithiothreitol (DTT) in ND96 for 5 min and subsequently transferred to a recording chamber containing standard Ringer’s solution [(in mM) 115 NaCl, 3 KCl, 1.8 CaCl2, 10 HEPES, pH 7.4]. Borosilicate electrodes (0.1–1 MΩ) filled with 3 M KCl were used for voltage recording and current injection. Cl- currents were acquired using a Warner OC-725C oocyte clamp (Harvard Apparatus, Holliston, MA, USA). Data were sampled at 10 kHz and filtered at 1 kHz. All recordings were conducted at room temperature (20-22 °C).

### RNA extraction and quantitative reverse transcription PCR (RT-qPCR)

Total RNA was extracted from transfected HEK293T cells using Quick-RNA Miniprep Kit (Zymo Research, Irvine, CA, USA) according to the manufacturer’s instructions. cDNA was synthesized from 1 μg of total RNA using the Magic RT cDNA synthesis kit (Bio-Genesis Technologies, Taiwan), following the manufacturer’s instructions. Quantitative PCR (qPCR) was conducted on a QuantStudio™ 5 Real-Time PCR Systems (Thermo Scientific) using TOOLS 2X SYBR qPCR Mix (Biotools, Xizhi Dist., New Taipei City, Taiwan). The specific primers used for PCRs are as follows: ClC-1 forward (5’-TGGAATCAGCGTGTTACGAG-3’) and reverse (5’-TCCCTATGTCCTGTCCTG TCCC-3’), GAPDH forward (5’-CAAGGTCATCCATGACAACTTTG-3’), and reverse (5’-GTCCACCACCCTGTTGCTGTAG-3’).

### Statistical analysis

Data are expressed as mean ± SD. Statistical significance between two groups was assessed using Student’s t-test, whereas comparisons among multiple groups were conducted with one-way ANOVA. All statistical analyses and curve fitting were carried out using Origin 7.0 (Microcal Software, Northampton, MA, USA) or Prism 6 (GraphPad Software, San Diego, CA, USA).

## RESULTS

### Myotonia congenita-associated ClC-1 alanine mutations

More than 350 mutations have been linked to myotonia congenita (Brenes et al., 2023). Among these disease-related *CLCN1* mutations, approximately 21 occur at alanine residues (Brenes et al., 2023; Suetterlin et al., 2022). Most of these mutations are located within the transmembrane domain and are distributed across eight different helices. Some of these myotonia-associated alanine mutations in the transmembrane domain involve substitutions with polar residues (*e.g.*, threonine), as well as non-polar residues (*e.g.*, valine). To further understand how myotonia-associated mutations involving alanine residues may cause significant structural or functional impairments of the ClC-1 channel, we decided to conduct biochemical analyses of some of these ClC-1 mutant proteins. We focused on the myotonia-causing ClC-1 mutations located at A218 (helix E), A313 (helix I), A493 (helix N), A531 (helix O), and A566 (helix Q) (Fig. 1A) (de Diego et al., 1999; Fialho et al., 2007; Kubisch et al., 1998; Mazon et al., 2012; Papponen et al., 1999; Skalova et al., 2013; Suetterlin et al., 2022).

We began by studying the effect of alanine-to-threonine mutations (A218T, A313T, A493T, A531T, A566T) on ClC-1 proteostasis. As shown in Figure 1B, total ClC-1 protein expression levels were substantially decreased for the helix O mutation A531T and helix Q mutation A566T, but not for their counterparts in helices E, I, and N. We also investigated the proteostatic impact of alanine-to-valine mutations (A218V, A313V, A493V, A531V, A566V) in ClC-1. Biochemical analysis revealed that these valine substitutions led to a notable decrease in expression levels of virtually all ClC-1 mutant proteins, except for the helix I mutation A313V (Fig. 1C). Taken together, our data suggest that the threonine/valine substitutions of alanine in helices O and Q appeared to result in more prominent disruption of ClC-1 proteostasis, with the helix O mutations A531T and A531V showing the most significant impact. This finding further implies that conformational changes at residue A531 may play a critical role in determining ClC-1 protein expression such that minor alterations of the side chain of this residue can considerably impair ClC-1 proteostasis.

### Effect of amino acid substitutions at residue 531 on ClC-1 protein expression

As highlighted by the effects of multiple helical alanine mutations shown in Figures 1B-C, compared to threonine substitutions, valine substitutions appeared to result in a more pronounced disruption of ClC-1 proteostasis. To address the potential mechanism underlying this differential impact of threonine/valine substitutions of helical alanine residues on ClC-1 proteostasis, we went on to generate a series of different mutations at A531 in helix O. Specifically, we introduced an additional 11 mutations at residue A531, including A531C, A531D, A531F, A531G, A531I, A531K, A531M, A531R, A531S, and A531Y. Figures 2A-B showed that, like the foregoing A531T and A531V mutations, all of these A531 substitutions led to a significant decrease in ClC-1 protein expression. For example, A531S, A531C, and A531V exhibited approximately 38%, 53%, and 80% reduction, respectively, in total protein levels.

**Figure 2.**
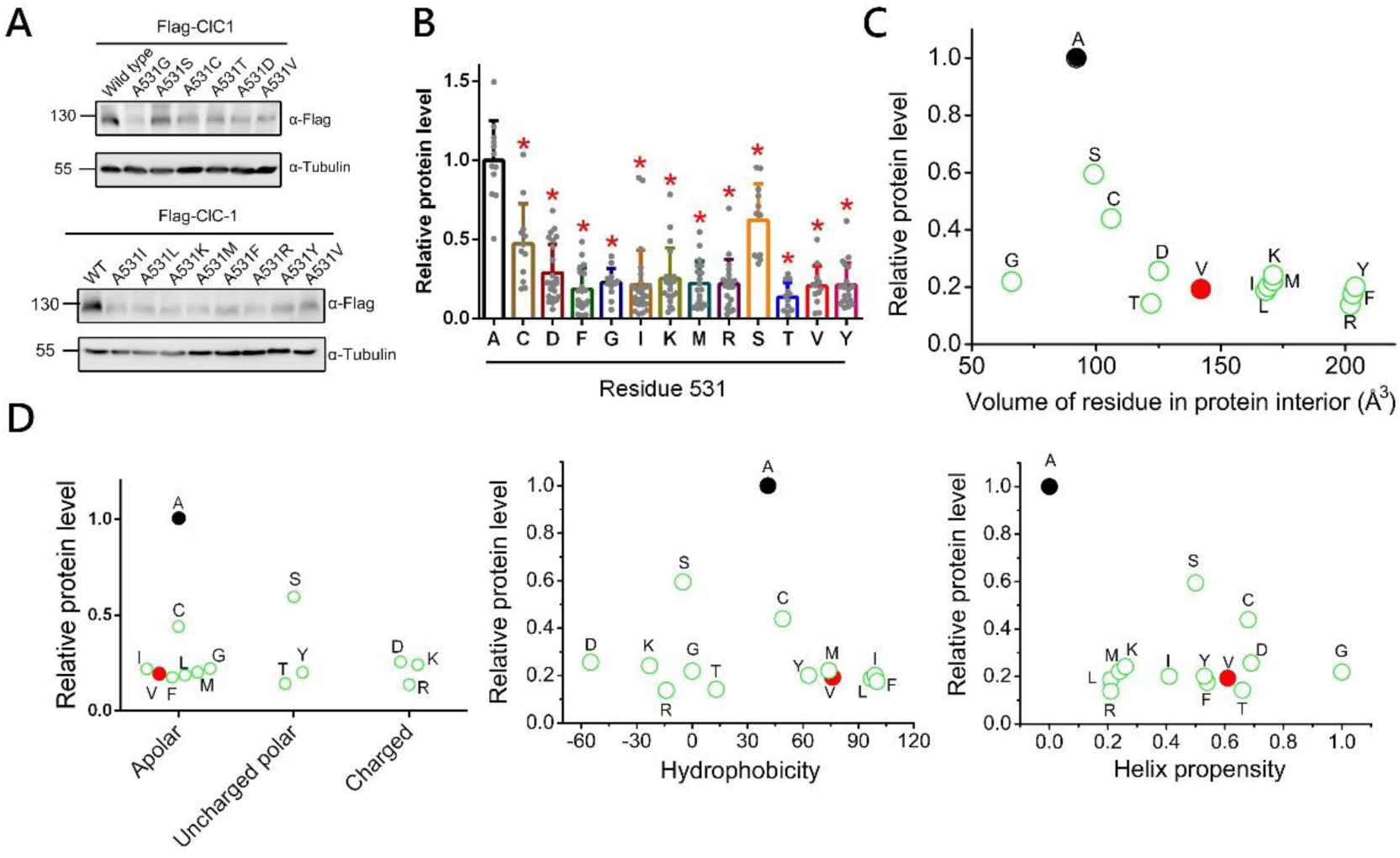
Effect of amino acid substitutions at residue 531 on ClC-1 protein expression. (A) Representative immunoblot of ClC-1 containing a series of mutations at residue A531. (B) Quantification of ClC-1 A531 substitution mutants. Normalized ClC-1 protein level: WT, 1.0±0.24; A531C, 0.47±0.25; A531D, 0.28±0.18; A531F, 0.18±0.13; A531G, 0.22±0.09; A531I, 0.21±0.19; A531K, 0.25±0.19; A531M, 0.22±0.14; A531R, 0.21±0.15; A531S, 0.62±0.22; A531T, 0.13±0.09; A531V, 0.20±0.12; A531Y, 0.21±0.14 (*, *P* < 0.05; n = 14–32). (C) ClC-1 total protein levels, normalized to wild type (alanine, A), plotted against side-chain volume. Wild type (alanine, A) is represented by a filled black circle; valine (V) as a filled red circle; and all other mutants as green open circles. (D) ClC-1 protein levels of A531 mutants normalized to wild type and plotted against (left) amino acid chemical structures (apolar, uncharged polar, charged), (middle) hydrophobicity, normalized to glycine (hydrophobicity index = 0), and (right) helix propensity, with alanine defined as 1.0 as the strongest helix-forming residue. Helix propensity values of other amino acids were derived from simulation.

One of the major impacts of these substitutions at residue 531 entails a change in the predicted volume of the amino acid side chain (Häckel et al., 1999; Harpaz et al., 1994; Zamyatnin, 1984). For example, substitution of alanine with serine results in about 6% increase in the residue volume (from 90 to 95.4 Å), while cysteine and valine substitutions lead to even larger increases by about 15% (from 90 to 103.3 Å) and 54.4% (from 90 to 139 Å), respectively (Tsai et al., 1999). We therefore decided to compare the relative ClC-1 protein expression level of these A531 substitution mutants based on the residue volume of the substituting amino acid. As shown in Figure 2C, increasing the residue volume at residue 531 appeared to progressively down-regulate ClC-1 protein expression. For instance, substitution with serine, cysteine and valine resulted in 41%, 57% and 81% reduction, respectively, in ClC-1 protein level. In contrast, no significant correlation was observed when we plotted relative ClC-1 protein expression level against other properties of the substituting amino acids, such as chemical structure (apolar, uncharged polar, charged), hydrophobicity (Monera et al., 1995), or helix propensity (Scholtz, 1998) (Fig. 2D). The only exception, however, concerns substitution with glycine, which, despite having the smallest residue volume, led to about 80% reduction in protein expression level, much more significant than most of the other mutants. This unexpected effect may be attributed to the unique structure of glycine, consisting of only a hydrogen as its side chain, thereby conferring low helical propensity and limiting its ability to stabilize α-helical structures (Javadpour et al., 1999; O’Neil & DeGrado, 1990).

To further verify this potential inverse relationship between ClC-1 protein level and the residue volume at residue 531, we went on to study the effect of A531 substitution on functional expression of ClC-1 channel. Electrophysiological analyses revealed that, as the residue volume at residue 531 increased, the corresponding current levels of the ClC-1 A531 substitution mutants decreased progressively (Fig. 3A; Suppl. Fig. S1). In contrast, no significant relationship was observed between the residue volume of the substituting amino acid and the voltage-dependent gating property of the ClC-1 mutant (Fig. 3B; Suppl. Fig. S2; Suppl. Table 1). Our further analysis demonstrated that this inverse relationship between the A531 residue volume and ClC-1 protein levels, as well as functional expression, could be well-fitted by a monoexponential decay equation, thereby quantifying the nature of this inverse correlation (Fig. 3C).

**Figure 3.**
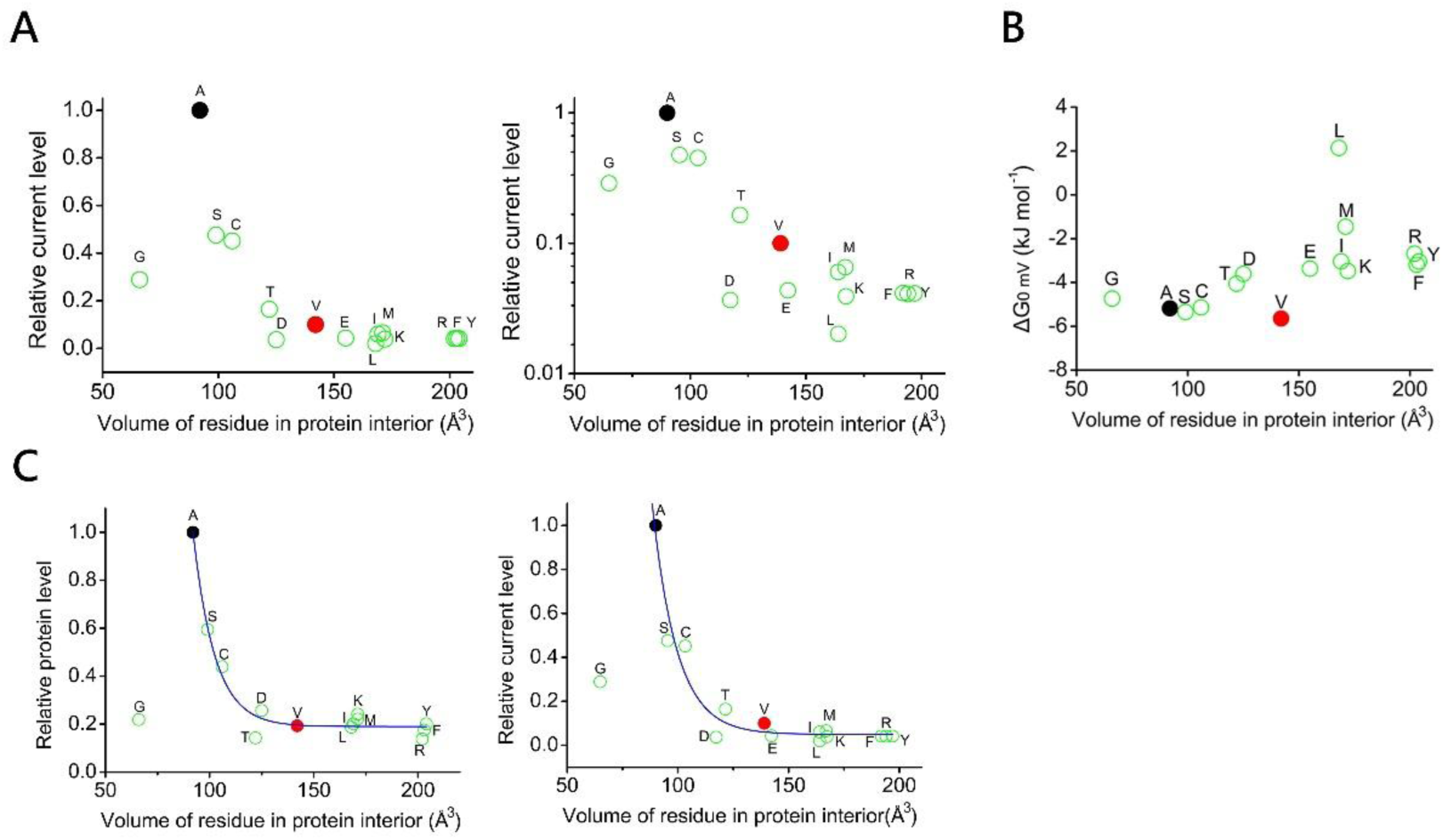
The inverse relationship between ClC-1 A531 residue volume and functional expression. (A) ClC-1 current levels of A531 mutants, normalized to WT, are plotted against residue volume, with the left panel shown on a linear scale and the right panel on a semi-logarithmic scale. Normalized ClC-1 current level: WT, 1.0±0.03; A531C, 0.45±0.02; A531D, 0.03±0.003; A531F, 0.04±0.01; A531G, 0.28±0.01; A531I, 0.06±0.02; A531K, 0.03±0.01; A531M, 0.06±0.02; A531R, 0.04±0.02; A531S, 0.47±0.02; A531T, 0.16±0.006; A531V, 0.10±0.02; A531E, 0.04±0.002; A531L, 0.02±0.01; A531Y, 0.04±0.01(*, *P* < 0.05; n = 22-133). (B) Free energy change (ΔG at 0 mV) of ClC-1 channels harboring A531 mutations, plotted as a function of residue volume. The free energy (ΔG_0_ mV) of the ClC-1 channel at 0 mV was derived using two formulas: ΔG_0_ mV = zF V_0.5_ and z = (RT/F) k^−1^. (C) Normalized (*left*) protein and (*right*) current levels of ClC-1 A531 mutants, plotted against side-chain volume and fitted with monoexponential equation Protein levels and current levels of A531 mutants are fitted with y= 5687.5 • e ^−x/10.4^ + 0.1 and y= 2940.7 • e ^−x/11.1^ + 0.04, respectively. A 63% reduction in ClC-1 protein and current levels is observed for every 10 Å^3^ and 11 Å^3^ increase in residue 531 volume, respectively.

Taken together, mutations at residue 531 in helix O lead to a pronounced reduction in total ClC-1 protein expression and current amplitude. The residue volume at this position appears to be a key determinant of ClC-1 channel protein, with alanine providing the optimal expression level.

### Disruption of ClC-1 protein stability by A531 substitutions

Attenuation of steady-state protein level may be caused by defective protein translation and/or reduced protein stability. RT-qPCR analysis revealed that mRNA levels were unchanged upon mutation at residue 531, indicating that these mutations do not affect ClC-1 transcription or translation (Suppl. Fig. S3). We therefore performed cycloheximide (CHX) chase assays to examine whether A531 substitutions disrupt the protein stability of ClC-1. As shown in Figure 4, ClC-1 protein half-life progressively decreased as the residue volume of the A531 substitutions increased. For example, ClC-1 A531 substitutions with larger residue volumes, including A531T, A531D, A531M, and A531K, reduced the protein half-life from 6.3 hours in the wild type (WT) to 2.4 hours, corresponding to a ∼62% decrease. Moreover, A531 substitutions with smaller residue volumes, such as A531S, showed a less pronounced reduction in protein half-life, decreasing from 6.3 hours to 4.4 hours, representing an approximate 30% reduction relative to the WT. Taken together, these results indicate that increasing the residue volume at residue A531 destabilizes ClC-1 protein and enhances its susceptibility to degradation, suggesting that the residue volume at this position plays a critical role in maintaining ClC-1 protein stability.

**Figure 4.**
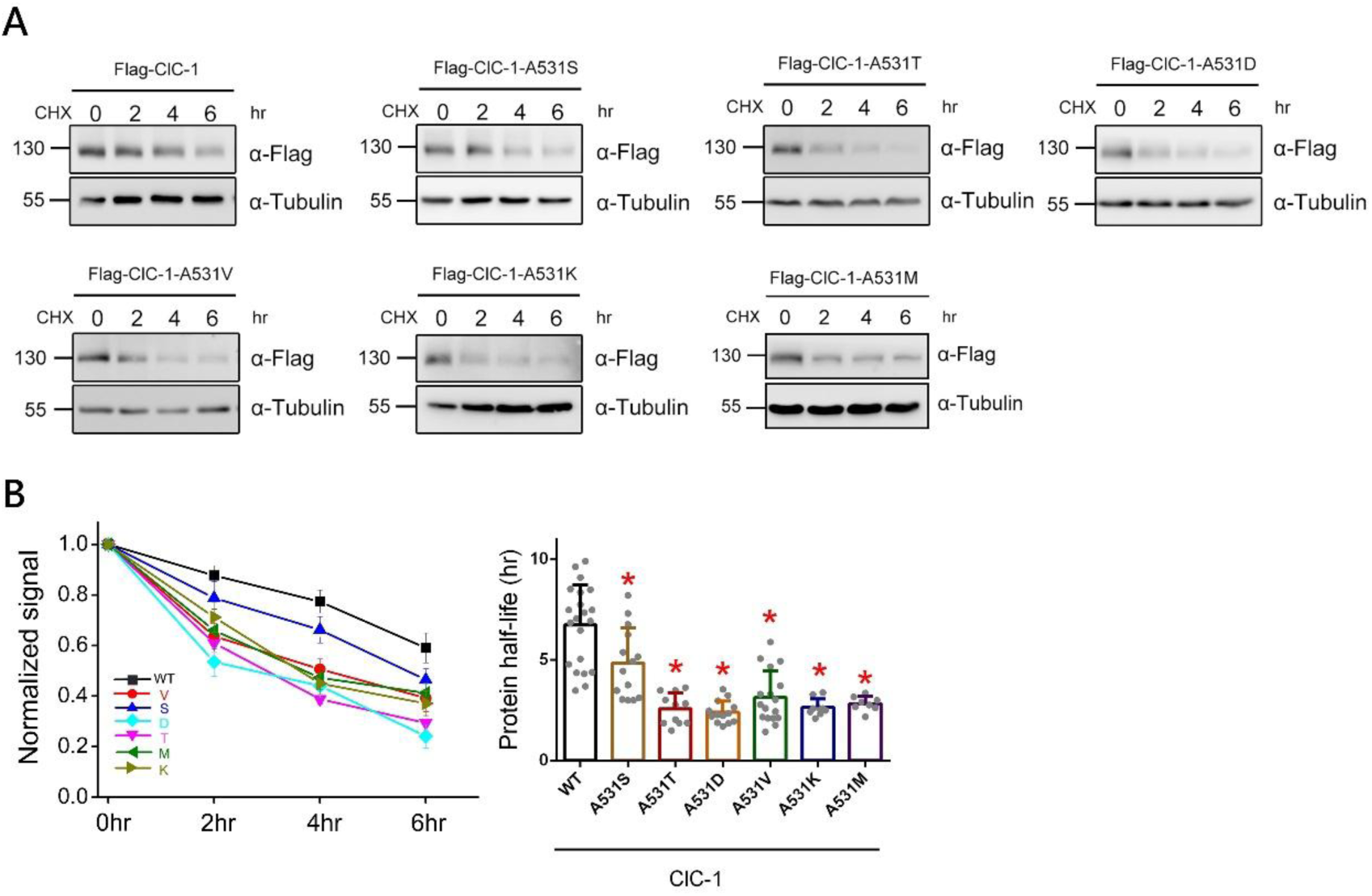
Disruption of ClC-1 protein stability by A531 substitutions. ClC-1 protein stability was altered by the introduction of A531 mutations. (A) Representative immunoblot showing the ClC-1 protein stability with various A531 mutations. Transfected HEK293T cells were subjected to 100 μg/ml cycloheximide (CHX) treatment for indicated durations (0-6h). (B) (*Left*) Linear plot showing the relative levels of ClC-1 protein following various durations of CHX treatment. Protein signals were quantified as the ratio of ClC-1 to the corresponding α-Tubulin signals, and subsequently normalized to the untreated control (0 h). (*Right*) Quantitative analysis of ClC-1 protein half-life in the presence of various A531 mutations. Normalized ClC-1 protein half-life: WT, 6.73±1.98; A531S, 4.84±1.74; A531T, 2.58±0.77; A531D, 2.41±0.55; A531V, 3.14±1.29; A531K, 2.65±0.42; A531M, 2.83±0.36 (*, *P* < 0.05; n = 11-22).

We previously demonstrated that the cullin 4 (CUL4)–damage-specific DNA binding protein 1 (DDB1)–cereblon (CRBN) E3 ubiquitin ligase complex mediated the endoplasmic reticulum–associated degradation (ERAD) of ClC-1 and ClC-2 channels (Chen et al., 2015; Fu et al., 2020). Building on this finding, we analyzed the interaction between mutant ClC-1 and CRBN to determine whether A531 substitutions influence the CUL4–DDB1–CRBN degradation pathway, thereby leading to ClC-1 destabilization. Through immunoprecipitation analysis, we observed that A531 substitutions with substantially larger residue volumes showed a significant increase in CRBN binding efficiency. Notably, A531M, which has the largest residue volume among the tested variants, exhibited a 5.5-fold increase in CRBN binding compared to the WT. Similarly, ClC-1 variants with A531T, A531D, A531V, or A531K substitutions displayed increased binding affinity for CRBN (Fig. 5A; Suppl. Fig. S4).

**Figure 5.**
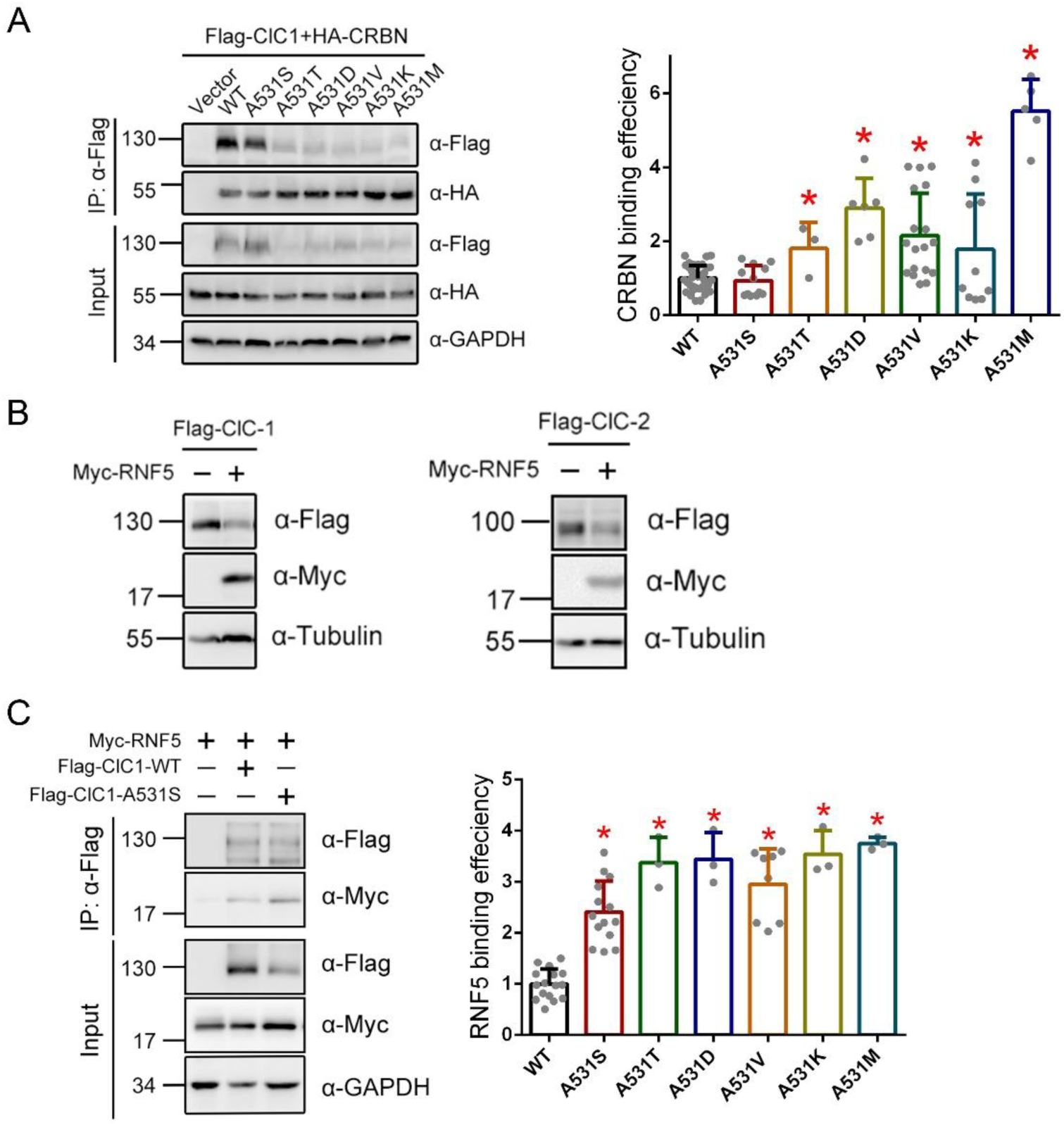
A531 substitutions enhance the interaction of ClC-1 with the substrate binding protein CRBN and E3 ubiquitin ligases RNF5. Co-immunoprecipitation of ClC-1 A531 mutations and HA-CRBN. (*Left*) Representative immunoblots. Co-expression with the Flag vector was used as the control. HEK293T cell lysates were immunoprecipitated (IP) with anti-Flag, followed by immunoblotting with anti-Flag or the anti-HA. Corresponding expression levels of ClC-1 and HA-CRBN in the lysates are shown in the “input” lane. Hereafter, input represents∼10%of the total protein used for immunoprecipitation. (*Right*) Quantitative analysis of CRBN binding efficiency for ClC-1 A531 substitutions. Binding efficiency was calculated as the ratio of input-normalized IP CRBN to input-normalized IP ClC-1. These ratios were subsequently normalized against the WT control to facilitate comparison across groups. Normalized ClC-1-CRBN binding efficiency: WT, 1.0±0.34; A531S, 0.93±0.40; A531T, 1.80±0.70; A531D, 2.89±0.80; A531V, 2.15±1.14; A531K, 1.78±1.49; A531M, 5.51±0.87 (*, *P* < 0.05; n = 3-36). (B) Representative immunoblot showing the effect of co-expression of Myc-RNF5 on (*Left*) ClC-1 or (*Right*) ClC-2 protein expression. (C) (*Left*) Representative immunoblots showing increased binding of ClC-1 A531S to RNF5 compared with the WT. (*Right*) Quantification of Myc-RNF5 binding efficiency for ClC-1-WT and A531 substitutions. To determine relative association, IP signals for Myc-RNF5 and Flag-ClC-1 were first normalized to their respective inputs. Data are presented as the ratio of RNF5/ClC-1 and normalized to WT. Normalized ClC-1-RNF5 binding efficiency: WT, 1.00 ± 0.29; A531S, 2.40±0.61; A531T, 3.37±0.49; A531D, 3.44±0.52; A531V, 2.95±0.69; A531K, 3.54±0.46; A531M, 3.75±0.12 (*, *P* < 0.05; n = 3-16).

In contrast, the A531S mutation, which represents only a minor increase in residue volume relative to alanine, exhibited a ClC-1–CRBN binding efficiency comparable to that of the WT (Fig. 5A). To further investigate the molecular mechanisms regulating ClC-1 A531S protein stability, we focused on the ER-resident E3 ligase RNF5, which has been reported to mediate early-stage ERAD of cystic fibrosis transmembrane conductance regulator (CFTR) and to modulate the stability of several membrane proteins (Li et al., 2023; Morito et al., 2008; Okiyoneda et al., 2018; Tsai et al., 2022). The co-expression of RNF5 notably reduced ClC-1 and ClC-2 protein levels, implying a regulatory role for RNF5 on ClC-1 and ClC-2 channels (Fig. 5B). As shown in Figure 5C, the ClC-1 A531S mutant displayed a pronounced increase in RNF5 binding efficiency compared with the WT. Consistently, all A531 substitutions significantly augmented the interaction between ClC-1 and RNF5. Collectively, these observations indicate that mutations at residue 531 promote ClC-1 degradation through the ERAD pathway and compromise ClC-1 protein stability mediated by the substrate binding protein CRBN and E3 ubiquitin ligase RNF5.

### Conserved alanine residues in helix O as determinants of CLC channel protein stability

Based on the cryoEM structure of ClC-1 (Park & MacKinnon, 2018) revealed that A562 in helix Q is the nearest alanine to A531 from another helical region, with a distance of about 3.5 Å. However, the substitution of A562 to threonine or valine failed to elicit a significant reduction in the ClC-1 protein expression level (Suppl. Fig. S5). These findings imply that the alanine substitutions within helix O may render the ClC-1 channel protein more susceptible to ERAD mechanisms. Moreover, sequence alignment of helix O further revealed the presence of five highly conserved alanine residues across the CLC channel/transporter superfamily (Fig. 6A). In ClC-1, the conserved alanine residues are A525, A529, A530, A531, and A535; in ClC-2, they are A500, A504, A505, A506 and A510. Importantly, ClC-1 A531V and ClC-2 A500V are associated with myotonia congenita and leukoencephalopathy, respectively (Depienne et al., 2013; Papponen et al., 1999). Consistent with the findings for A531 substitutions, replacement of these highly conserved alanine residues in helix O with valine resulted in a significant reduction in total protein expression levels of both ClC-1 and ClC-2 (Fig. 6B-C). Altogether, these data underscore the critical role of helix O in CLC channel protein stability, particularly the contribution of highly conserved alanine residues. Substitution of these alanine residues disrupts CLC protein stability, indicating that alanine provides the most favorable condition for maintaining proper CLC proteostasis.

**Figure 6.**
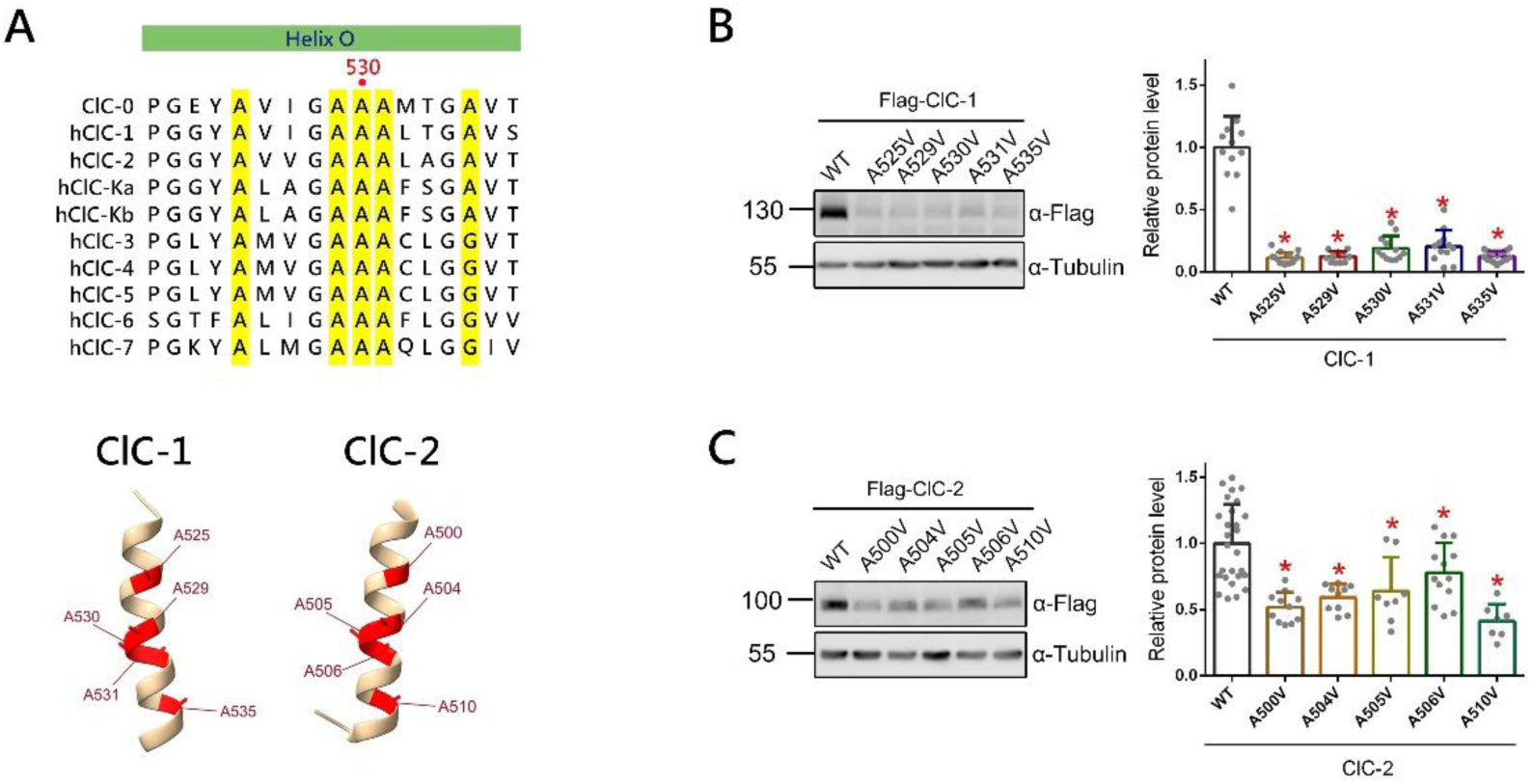
Conserved alanine residues in helix O as determinants of CLC channel protein stability. (A) (*Upper*) Protein sequence alignment of the helix O region of human CLC family members and ClC-0 from *Torpedo*. Conserved alanine residues across the CLC family are highlighted in yellow. (*Bottom*) Structures of human ClC-1 helix O (PDB: 6COY) and ClC-2 helix O (PDB: 7XF5). Conserved alanine residues in both structures are highlighted in red. (B) Immunoblot analysis and quantification of ClC-1 wild type and mutants in which conserved alanine residues were substituted with valine. Normalized ClC-1 protein level: WT, 1.0±0.24; A525V, 0.11±0.04; A529V, 0.12±0.04; A530V, 0.18±0.09; A531V, 0.20±0.12; A535V, 0.12±0.04 (*, *P* < 0.05; n = 11-18). (C) Immunoblot analysis and quantification of ClC-2 wild type and mutants in which conserved alanine residues were substituted with valine. Normalized ClC-2 protein level: WT, 1.0±0.29; A500V, 0.51±0.11; A504V, 0.59±0.10; A505V, 0.63±0.25; A506V, 0.77±0.22; A510V, 0.41±0.12 (*, *P* < 0.05; n = 7-27).

### Adjusting the spatial arrangement of residues surrounding ClC-2 A500V can restore impaired protein stability

As illustrated in Figures 6B-C, the disease causing ClC-2-A500V mutation exhibited a less pronounced reduction in protein level compared to the disease related ClC-1-A531V mutant. We therefore focused our investigation on the ClC-2-A500V mutation to explore potential strategies for rescuing its defective protein stability. Notably, the increased residue volume from alanine residue in helix O is likely to reduce the interhelical spacing between helix O and its adjacent helices. We therefore investigated whether structurally expanding the space adjacent to the mutation facilitates a necessary rearrangement of ClC-2-A500V and its neighboring helices, thereby restoring proteostasis. Using UCSF ChimeraX and the PDB structure 7XF5, we identified ClC-2 residues A467 and R471, which are located in close spatial proximity to the disease-associated residue A500V. As shown in Figure 7A, the distance between ClC-2 A500V and the nearby residue R471 is approximately 1.69 Å, which increased to 3.31 Å upon substitution of R471 with alanine (R471A), a smaller side-chain residue. In addition, the distance between A500V and its neighboring residue A467 is 2.47 Å. Upon introduction of the A467G mutation, the distance could not be precisely measured due to the absence of a side chain in glycine; however, the structural model suggests an expansion of the inter-residue spacing (Fig. 7B).

**Figure 7.**
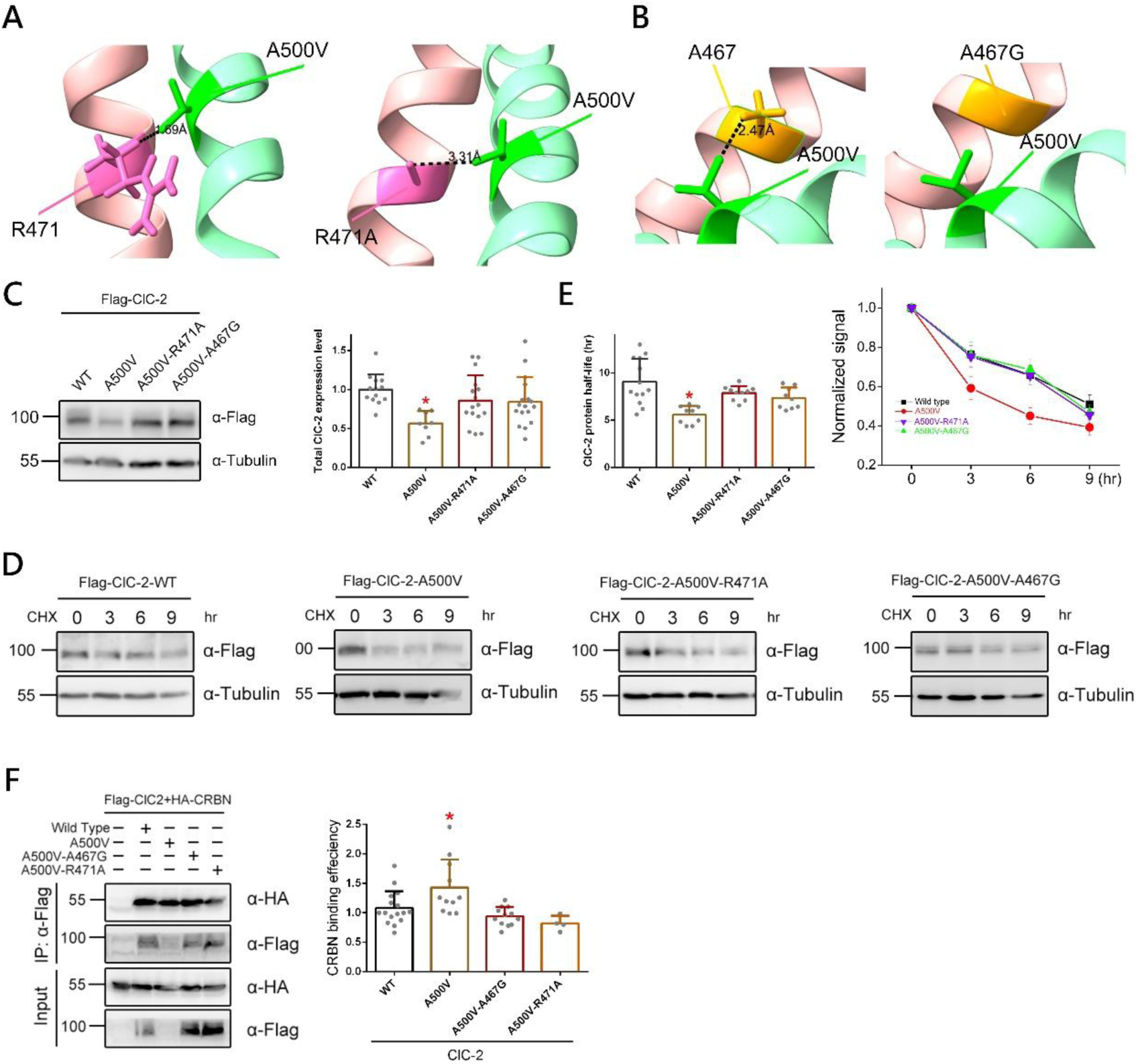
Adjusting the spatial arrangement of helices surrounding helix O can restore impaired protein stability. (A-B) Modeling of the side-chain spatial configuration of the ClC-2 A500V mutant and the nearby residue R471 and A467 (PDB: 7XF5). A500V is shown in dark green within helix O (light green), while R471 and A467 are depicted in dark pink and orangr, respectively, within helix N (pale pink). (*left*) original structure of the A500V mutant with residues R471 and A467. (*right*) spatial configuration of the R471A-A500V and A467G-A500V double mutant. The distance between the two residues is indicated by a black dashed line. (C) Representative immunoblots and quantification of ClC-2 A500V mutant compare to double mutants. Normalized ClC-2 protein level: WT, 1.0±0.19; A500V, 0.56±0.15; A500V-R471A, 0.85±0.32; A500V-A467G, 0.84±0.31 (*, *P* < 0.05; n = 8-17). (D) Representative immunoblot showing the ClC-2 double mutants applying cycloheximide (CHX) assay. Transfected HEK293T cells were subjected to 100 μg/ml cycloheximide (CHX) treatment for indicated durations (0-9h). (E) (*left*) Quantitative analysis of ClC-2 protein half-life in the presence of A500V or other double mutations. Normalized ClC-2 protein half-life: WT, 9.07±2.40; A500V, 5.57±0.93; A500V-R471A, 7.86±0.74; A500V-A467G, 7.32±1.12 (*, *P* < 0.05; n = 9-14). (*right*) Linear plot showing the relative levels of ClC-2 protein following various durations of CHX treatment. Protein signals were quantified as the ratio of ClC-2 to the corresponding *α*-Tubulin signals, and subsequently normalized to the untreated control (0 h). (F) Representative immunoblots of co-immunoprecipitation between ClC-2 mutants and HA-CRBN. The right panel shows quantitative analysis of CRBN binding efficiency for ClC-2 A500V and double mutants. HA-CRBN signals were normalized to Flag-ClC-2 signals to calculate the binding efficiency ratio. Normalized ClC-2-CRBN binding efficiency: WT, 1.0±0.28; A500V, 1.43±0.47; A500V-A467G, 0.94±0.15; A500V-R471A, 0.82±0.12 (*, *P* < 0.05; n = 9-16).

To further investigate the effect of restoring the spatial arrangement surrounding helix O on ClC-2 proteostasis, total ClC-2 protein expression was examined. Figure 7C shows that introducing additional mutations, R471A or A467G, significantly restored the reduced total protein expression of ClC-2 A500V. Moreover, CHX chase analysis indicated that the protein half-life of ClC-2 A500V-R471A and A500V-A467G were significantly increased compared to ClC-2 A500V, and were restored to WT levels (Fig. 7D-E). To further elucidate the influence of helix O spatial rearrangement on ClC-2 proteostasis during ER quality control, we examined the binding efficiency of these mutants with CRBN. Co-immunoprecipitation analysis revealed that ClC-2 proteins harboring double mutations exhibited significantly reduced CRBN binding efficiency compared to ClC-2 A500V (Fig. 7F). Conversely, the proposed interhelical spatial rearrangement surrounding ClC-1 A531 did not succeed in restoring the protein stability of the myotonia associated ClC-1 A531V mutant (Suppl. Fig. S6).

Altogether, these findings suggest that reorganization of helix O toward a WT–like conformation reduces ER-associated ubiquitination and degradation, thereby restoring ClC-2 protein stability.

### Alanine mutations in helix O disrupt CLC channel trafficking

Given that CLC channels exert their physiological functions at the plasma membrane, it is essential to assess whether spatially rearranged channels are properly trafficked to the membrane and retain functional activity. However, subsequent electrophysiology recordings demonstrated that the ClC-2 double mutants failed to restore the ClC-2 current level, which remained undetectable and similar to the A500V single mutant (Fig. 8A). Surface biotinylation analysis further revealed that the ClC-2 double mutant protein expression levels at the cell surface were significantly decreased compared to the WT, remaining comparable to the A500V single mutant (Fig. 7B). To further examine the effect of ClC-2 helix O alanine mutations at the plasma membrane, ClC-2 channel helix O alanine substitutions were subjected to further analysis. Both electrophysiological recordings and biotinylation assays suggested that the helix O alanine mutations resulted in a markedly decreased ClC-2 current level and ClC-2 surface expression level (Fig. 8C-D; Suppl. Fig. S7).

**Figure 8.**
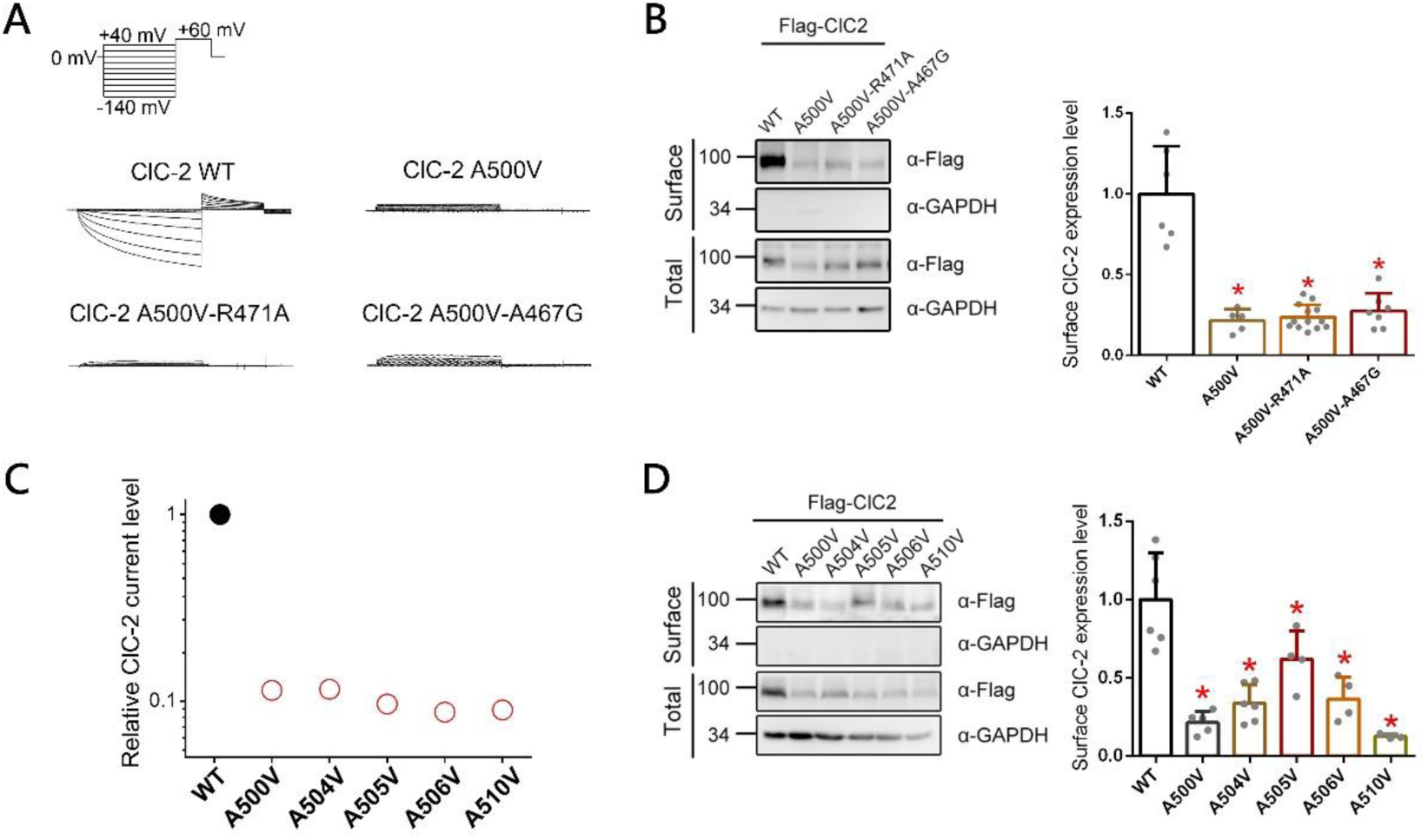
Alanine mutations disrupt ClC-2 channel trafficking. (A) Representative Cl^−^ current traces recorded via whole cell patch clamp in HEK293T cells expressing ClC-2-WT, A500V, A500V-R471A and A500V-A467G. From a holding potential of 0 mV, cells were subjected to 1 s voltage steps ranging from –140 to +40 mV in 20 mV increments, followed by a tail-voltage step to +60mV for 500 ms, as shown in the left panel. Cells expressing ClC-2-WT were used as controls. (B) Surface biotinylation analysis showing that ClC-2 double mutants failed to restore cell-surface protein levels compared with the ClC-2 A500V mutant. (*left*) Representative immunoblots of biotinylated HEK293T cell lysates. Total protein was analyzed directly, whereas surface protein was enriched by streptavidin pull-down prior to immunoblotting. GAPDH was included as a loading control. (*right*) Quantification of surface protein expression. Surface protein intensities were first standardized to the corresponding total ClC-2 signals and subsequently normalized to the WT control. Normalized surface ClC-2 protein level: WT, 1.0±0.29; A500V, 0.21±0.07; A500V-R471A, 0.23±0.07; A500V-A467G, 0.27±0.10 (*, *P* < 0.05; n = 5-13). (C) Relative ClC-2 current levels recorded in Xenopus oocytes, normalized to their corresponding WT controls. The Y-axis is shown on a semi-logarithmic scale. (D) Mutations at conserved alanine residues in helix O exhibit decreased ClC-2 surface expression level. (*left*) Representative immunoblots. (*right*) Quantification of surface protein expression. Surface protein intensities were first standardized to the corresponding total ClC-2 signals and subsequently normalized to the WT control. Normalized surface ClC-2 protein level: WT, 1.0±0.29; A500V, 0.21±0.07; A504V, 0.33±0.11; A505V, 0.61±0.18; A506V, 0.36±0.13; A510V, 0.12±0.01(*, *P* < 0.05; n = 3-6).

In addition to ClC-2, the effect of helix O alanine substitutions on ClC-1 surface expression was subsequently investigated. As shown in Figure 9A, the substitutions at residue 531 not only disrupted ClC-1 protein stability, as previously mentioned, but also significantly decreased the ClC-1 surface expression level. Furthermore, all of the alanine mutations introduced into helix O resulted in a concurrent decrease in both the ClC-1 surface protein expression level and the corresponding current level (Fig. 9B; Suppl. Fig. S8). Since the helix O alanine residues were substituted with valine, which has a relatively larger residue volume, we examined the effect of residue volume on ClC-1. As shown in Figures 9C and 9D, both alanine-to-threonine and alanine-to-valine mutations reduced surface expression, with the A-to-T mutants showing a milder decrease relative to the A-to-V mutants.

**Figure 9.**
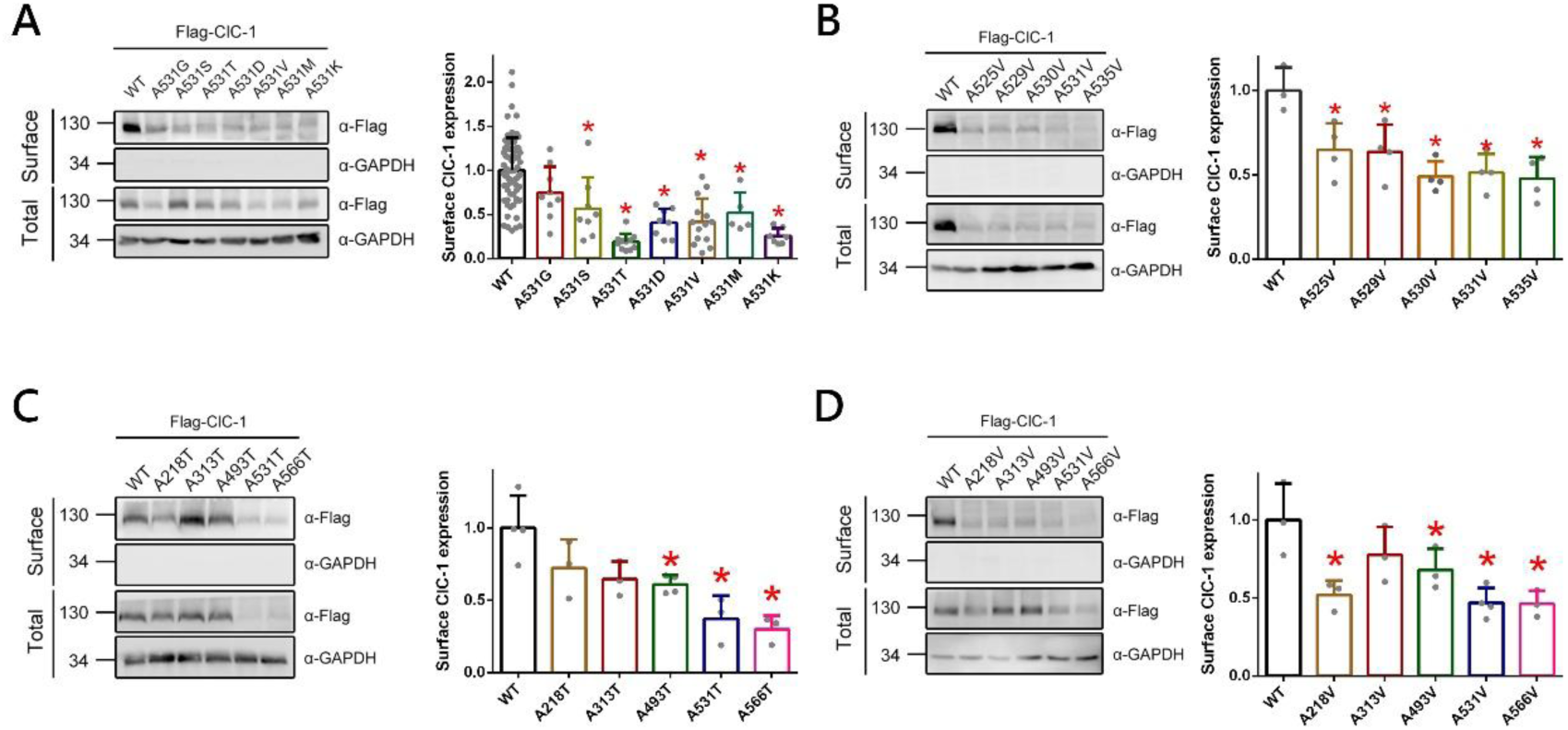
Alanine residue mutations exacerbate defects in ClC-1 protein trafficking. Representative immunoblots and quantitative analysis of biotinylated HEK293T cell lysates expressing (A) ClC-1 A531 substitutions, (B) mutations at conserved alanine residues in ClC-1 helix O, (C) alanine-to-threonine myotonia-associated mutants, and (D) alanine-to-valine myotonia-associated mutants. Total protein was analyzed directly, whereas surface protein was enriched by streptavidin pull-down prior to immunoblotting. GAPDH was included as a loading control. Surface protein intensities were first standardized to the corresponding total ClC-1 signals and subsequently normalized to the WT control. Normalized surface ClC-1 protein level: (A) WT, 1.0±0.25; A531G, 0.62±0.20; A531S, 0.50±0.16; A531T, 0.34±0.10; A531D, 0.42±0.11; A531V, 0.29±0.10; A531M, 0.13±0.04; A531K, 0.33±0.09 (B) WT, 1.0±0.06; A525V, 0.38±0.01; A529V, 0.26±0.02; A530V, 0.16±0.02; A531V, 0.14±0.006; A535V, 0.14±0.02 (C) WT, 1.0±0.22; A218T, 0.73±0.13; A313T, 0.80±0.20; A493T, 0.90±0.20; A531T, 0.34±0.10; A566T, 0.25±0.09 (D) WT, 1.0±0.21; A218V, 0.29±0.10; A313V, 0.47±0.11; A493V, 0.52±0.14; A531V, 0.18±0.05; A566V, 0.16±0.02 (*, *P* < 0.05; n = 3-12).

Taken as a whole, these data suggest that residue volume is among the factors influencing CLC channel trafficking. Moreover, although spatial rearrangement can restore the impaired protein stability caused by mutations that expand residue residue volume, it fails to rescue surface expression, implying that additional determinants beyond spatial conformation also contribute to CLC channel trafficking.

## DISCUSSION

Herein, we demonstrated the critical role of helix O, particularly its highly conserved alanine residues, in ClC-1 and ClC-2 proteostasis. The data suggest that substituting these alanine residues with other amino acids increases the susceptibility of CLC channels to degradation via the ER associated degradation pathway. As illustrated in the schematic model (Fig. 10A), helix O in wild type ClC-1 and ClC-2 exhibits a high conservation of alanine residues. Under functional, properly folded states, most ClC-1 and ClC-2 channels exit the ER and traffic to the plasma membrane to execute their functions, leaving only a small population to be targeted by RNF5 or CUL4–DDB1–CRBN E3 ubiquitin ligase complexes for proteasomal degradation. When mutations occur at these conserved alanine residues within helix O, a substantial proportion of the mutant CLC channels are recognized by the RNF5 and CUL4–DDB1–CRBN E3 ubiquitin ligase complexes for degradation via the ERAD pathway. Consequently, only a small fraction of mutant channels manage to traffic to the plasma membrane and exert their physiological functions (Fig. 10B).

**Figure 10.**
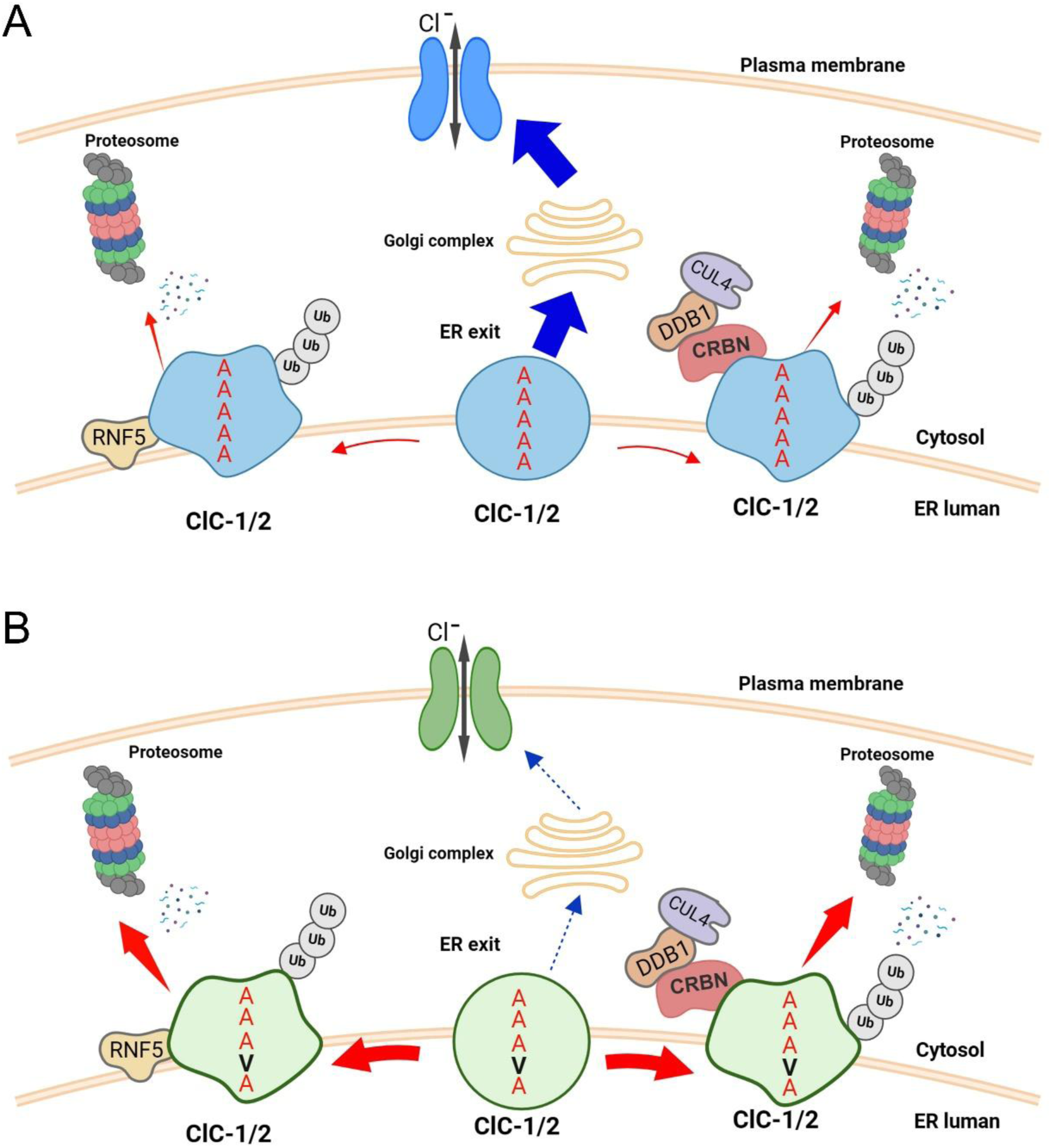
Schematic model illustrating the role of the highly conserved alanine residues within helix O in regulating CLC channel stability via the ER associated degradation pathway. (A) Under normal physiological conditions, correctly folded ClC-1 and ClC-2 channels, which carry five conserved alanine residues in helix O (highlighted in red), exit the ER and undergo trafficking to their final destination at the plasma membrane, where they exert their physiological functions. A small fraction of ClC channels is recognized by the ER-resident E3 ligase RNF5 and the CUL4–DDB1–CRBN E3 ubiquitin ligase complex, targeting them for degradation. (B) Mutations at conserved alanine residues within helix O alter the structure and conformation of ClC-1 and ClC-2, thereby leading to their recognition by the ER associated degradation machinery. Misfolded CLC channel are recognized at an early stage of ERAD by the ER-resident E3 ligase RNF5, which ubiquitinates and directs them to proteasomal degradation. ER-associated degradation of misfolded CLC channels is also mediated by the CUL4–DDB1–CRBN E3 ubiquitin ligase complex, which facilitates their ubiquitination and subsequent proteasomal degradation. Improper folding of CLC channel proteins results in defective membrane trafficking; consequently, only a small fraction of CLC proteins reaches the plasma membrane to exert their physiological functions. Created with BioRender.com.

A notable subset of disease-associated mutations in CLC channels that disrupt proteostasis involves the substitution of alanine residues with other amino acids. (Depienne et al., 2013; Fialho et al., 2007; Mazon et al., 2012; Papponen et al., 2008; Skalova et al., 2013). These findings highlight the critical role of alanine residues in CLC channel proteins, where their disruption is sufficient to impair channel function and contribute to disease pathogenesis. Notably, our analyses revealed that among the myotonia-associated A-to-T and A-to-V mutations, substitutions at residue A531 in helix O confer the most severe impairment of ClC-1 protein stability (Fig. 1B-C). Moreover, alterations of the highly conserved alanine residues in helix O markedly decreased the stability of both ClC-1 and ClC-2, whereas substitutions of alanine residues in helix Q did not uniformly compromise protein stability. Specifically, substitutions at A566 disrupted ClC-1 stability, whereas those at A562 had no detectable effect (Fig. 1B-C; Suppl. Fig. S6). Previous study revealed that in CLC exchangers, movements of helix O are transduced to the ion pathway through direct contact with its C-terminus (Basilio et al., 2014). In ClC-1 channel, myotonic mutant (G523D) at helix O showed a reversed voltage dependency comparing to WT (Seong et al., 2017). Taken together, these results underscore the significance of helix O, particularly its highly conserved alanine residues, as a critical determinant of ClC-1 and ClC-2 protein stability.

To further explore the relationship between the conserved alanine residues in helix O and ClC-1 protein stability, our findings indicate that mutations with larger residue volumes, such as valine, exhibited more pronounced defects in protein stability compared to those with smaller side chains, for instance threonine. This directly suggests a correlation between the substituting amino acid’s residue volume and ClC-1 proteostasis. Furthermore, functional expression analyses demonstrated that the current level of ClC-1 A531 substitutions exhibited an inverse correlation with the residue volume at residue 531, whereas the channels gating properties remained unaltered by these substitutions (Fig. 2; Suppl. Fig. S2). These findings are highly consistent with our previous functional expression studies, which showed that the A531V mutant specifically exhibited a significant reduction in ClC-1 current level while retaining largely unchanged gating properties. (Lee et al., 2013; Peng et al., 2016). Collectively, these data robustly establish that A531 substitutions in helix O predominantly compromise ClC-1 protein stability rather than inducing functional channel disruption. Critically, the degree of stability impairment follows a residue volume dependent manner, with increasing side chain bulk leading to a more profound defect

We have previously demonstrated that the reduced protein expression level of ClC-1 A531V can be rescued by the application of proteasome inhibitor-MG132 (Lee et al., 2013). In addition, treatment with the heat shock protein 90 (Hsp90) inhibitor 17-AAG significantly increased the protein expression level of the leukodystrophy-associated ClC-2 A500V mutant (Fu et al., 2021). These data indicate that the diminished protein expression levels of mutant ClC-1 result from enhanced degradation of misfolded proteins. Further biochemical analyses demonstrated that A531 substitutions in ClC-1 promoted interactions with the E3 ligases CRBN and RNF5, thereby accelerating ER quality control–mediated degradation and impairing ClC-1 protein stability.

Notably, RNF5 has been demonstrated to recognize misfolded substrates co translationally, allowing it to monitor the folding status of its targets at the ER membrane during the early stages of protein biogenesis (Morito et al., 2008; Younger et al., 2006). Interestingly, our results demonstrated that the A531S substitution, which introduces a smaller side chain, did not enhance binding to CRBN, while its interaction with RNF5 was markedly increased. These data suggest that RNF5 may recognize misfolded ClC-1 at the ER membrane prior to CRBN. Consequently, the relatively minor structural alteration induced by the ClC-1 A531S mutant would be more susceptible to RNF5 targeting and subsequent degradation through the ER quality control pathway

Our data revealed that the impaired ClC-2 protein stability resulting from the leukodystrophy related ClC-2 A500V mutant could be rescued through the rearrangement of the surrounding interhelical space, as evidenced by improvements in both the protein expression level and CRBN binding efficiency (Fig. 6). However, the protein stability of the myotonia associated ClC-1 A531V mutant failed to improve following the proposed interhelical spatial rearrangement surrounding residue 531 (Suppl. Fig. S6). It remains to be determined whether a more optimal rearrangement of the structural environment surrounding ClC-1 A531 could restore protein stability.

Although the spatial modification successfully restored the protein stability of the ClC-2 A500V mutation and prevented its degradation via ER quality control, it failed to rescue the ClC-2 surface expression level (Fig. 7). Similarly, several myotonia-associated alanine substitutions, despite displaying total protein expression levels comparable to the WT, demonstrated defective surface expression of ClC-1 (Fig. 8). The potential reason for this defective CLC channel surface expression level could be that, aside from ER-ssociated degradation, a separate peripheral quality control mechanism regulates protein stability directly at the plasma membrane. Collectively, these data suggest that CLC channel trafficking is highly sensitive to subtle structural alterations. However, the detailed mechanism by which these alterations impair CLC channel trafficking remains to be elucidated.

In summary, this study establishes helix O as a pivotal determinant of ClC-1 and ClC-2 channel proteostasis. The highly conserved alanine residues within this region are essential for stability, and even subtle substitutions destabilize the channels, underscoring alanine as the optimal residue for sustaining CLC channel homeostasis.

## ACKNOWLEDGEMENTS

We thank the staff of the Biomedical Resource Core at the First Core Labs, National Taiwan University College of Medicine, for excellent technical assistance in plasmid construction.

## ABBREVIATIONS

CBS: cystathionine-β-synthase
CFTR: cystic fibrosis transmembrane conductance regulator
CHX: cycloheximide
CRBN: cereblon
CUL4: cullin 4
DDB1: DNA damage-binding protein 1
DMEM: Dulbecco’s modified Eagle’s medium
DTT: dithiothreitol
E3: E3 ubiquitin ligase
ER: endoplasmic reticulum
ERAD: endoplasmic reticulum–associated degradation
HEK: human embryonic kidney
Hsp90: heat shock protein 90
IP: immunoprecipitation
PBS: phosphate buffered saline
PDL: poly-D-lysine
PMSF: phenylmethylsulfonyl fluoride
Po: open probability
RNF5: RING-finger protein 5
RT-qPCR: quantitative reverse transcription polymerase chain reaction
TEVC: two-electrode voltage clamp
Ub: ubiquitin
WT: wild-type

