## Supplemental files for "Proteostatic significance of helix O alanine residues in CLC channels"

### SUPPLEMENTARY TABLE

| | | $V_{0.5}$ (mV) | *k* | $\Delta G_{0\,mV}$ (kJ mol$^{-1}$) | n |
|---|---|---|---|---|---|
| Cell-attached | WT | -16.7 ± 3.5 | - | 36.2 ± 1.3 | 19 |
| | A531G | -18.3 ± 4.2 | - | 40.9 ± 3.1 | 13 |
| | A531S | -17.3 ± 4.7 | - | 46.0 ± 3.4 | 17 |
| | A531C | -10.7 ± 5.2 | - | 48.7 ± 5.3 | 30 |
| | A531T | -12.8 ± 1.5 | | 49.1 ± 3.6* | 17 |
| Whole-cell | WT | -83.1 ± 1.2 | -5.18 ± 0.46 | 39.5 ± 2.7 | 14 |
| | A531V | -85.5 ± 3.3 | -5.64 ± 0.47 | 37.3 ± 2.6 | 13 |
| | A531G | -82.0 ± 1.9 | -4.74 ± 0.17 | 42.6 ± 1.2 | 19 |
| | A531S | -91.9 ± 1.3 | -5.35 ± 0.20 | 42.3 ± 1.5 | 11 |
| | A531C | -91.6 ± 2.3 | -5.14 ± 0.28 | 43.9 ± 2.0 | 12 |
| | A531T | -88.3 ± 1.3 | -4.05 ± 0.26* | 53.6 ± 2.9* | 18 |
| | A531L# | 42.8 ± 2.9* | 2.14 ± 0.17* | 49.2 ± 1.5* | 17 |
| | A531I | -55.9 ± 2.6* | -3.04 ± 0.12* | 45.3 ± 3.2* | 12 |
| | A531M | -26.3 ± 3.9* | -1.45 ± 0.25* | 44.5 ± 2.5* | 12 |
| | A531F | -71.4 ± 2.4* | -3.19 ± 0.11* | 51.1 ± 1.9* | 8 |
| | A531Y | -66.2 ± 2.7* | -3.05 ± 0.10* | 53.5 ± 2.4* | 10 |
| | A531D | -64.7 ± 3.8* | -3.61 ± 0.23* | 44.1 ± 1.8* | 16 |
| | A531E | -61.4 ± 2.4* | -3.37 ± 0.22* | 44.9 ± 2.0* | 15 |
| | A531K | -64.5 ± 2.6* | -3.48 ± 0.15* | 45.6 ± 2.3* | 16 |
| | A531R | -62.0 ± 2.2* | -2.69 ± 0.18* | 56.8 ± 2.5* | 12 |

***Supplementary Table S1. The V0.5 values and Po–V gating properties of ClC-1 WT and A531 substitutions.***

Po–V curves were obtained from HEK293T cells expressing ClC-1. $V_{0.5}$ indicates the voltage at which half of the channels are open, k denotes the slope factor, and n represents the number of recorded cells.

### SUPPLEMENTARY FIGURES

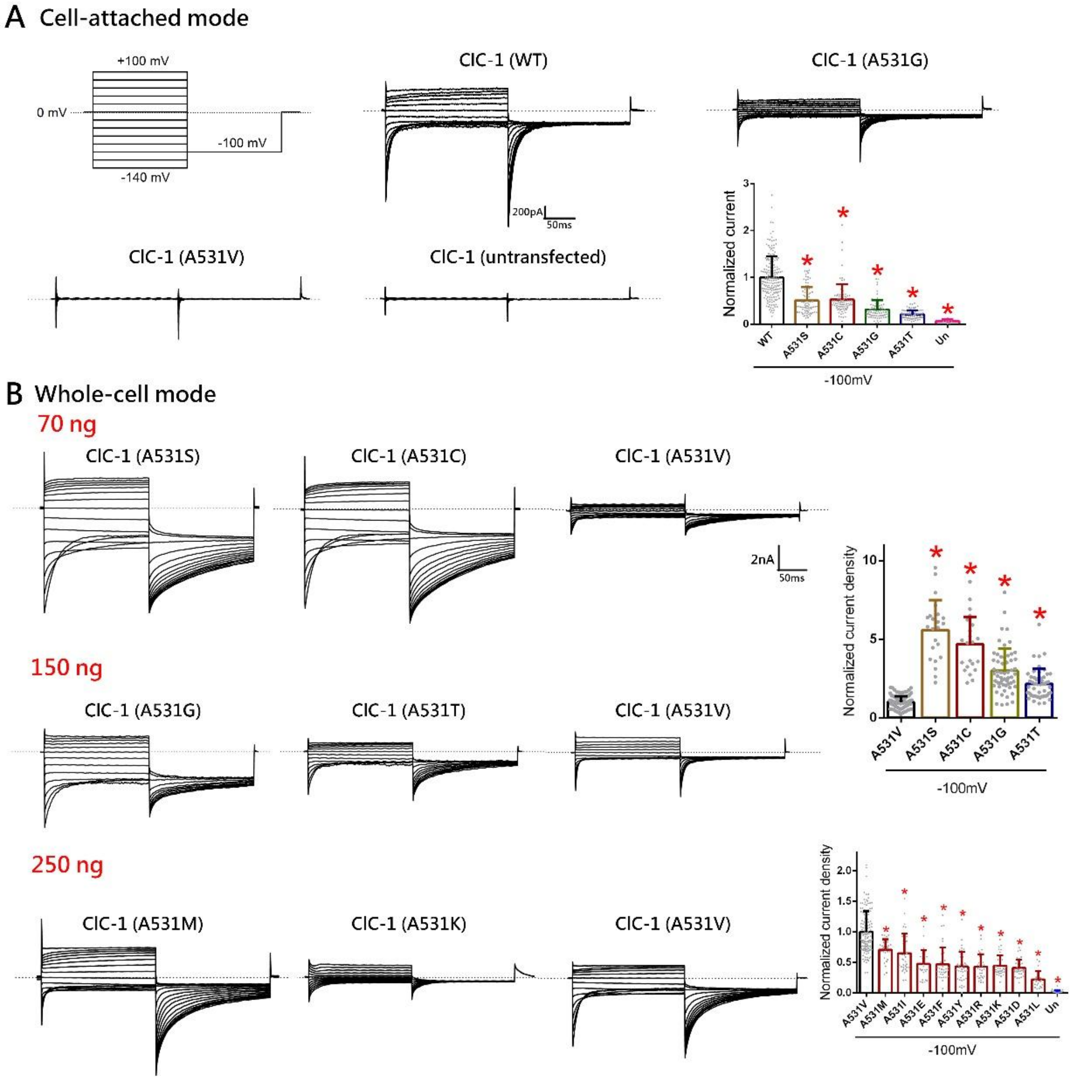


***Supplementary Figure S1. A531 substitutions significantly reduced ClC-1 current level compare to WT***

(A) Cell-attached patch-clamp recordings in HEK293T cells expressing ClC-1 WT, ClC-1 A531S, ClC-1 A531C, ClC-1 A531G, ClC-1 A531T, and in untransfected (un) control cells. (*Left*) Representative $Cl^-$ current traces recorded from human ClC-1 channels. From a holding potential of 0 mV, cells were subjected to 200-ms voltage steps ranging from +100 to –140 mV in –20 mV increments, followed by a tail-voltage step to -100 mV for 200 ms, as shown in the upper-left corner. (*Bottom-right*) Quantification of relative peak $Cl^-$ current amplitude at -100 mV. Data were normalized with respect to the ClC-1-WT control. Normalized ClC-1 current amplitude: WT, 1.00±0.44; A531C, 0.50±0.28; A531C, 0.49±0.32; A531G, 0.31±0.19; A53T, 0.20±0.09; untransfected, 0.06±0.03. Asterisks denote significant difference from the vector control (*, $P < 0.05$; n = 10-183) (B) Whole-cell patch-clamp experiments performed in HEK293T cells expressing different concentration of ClC-1 cDNA. (*Left*) Representative $Cl^-$ current traces recorded from human ClC-1 channels. cDNA concentrations used for transfection were 70 ng for ClC-1 A531S, A531C, and A531V; 150 ng for ClC-1 A531G, A531T, and A531V; and 250 ng for ClC-1 A531M, A531K, and A531V. Cells expressing ClC-1 with the A531V were used as controls. (*Right*) Quantification of relative peak $Cl^-$ current density at -100 mV. Data were normalized with respect to the A531V coexpressing control. The upper panel shows recordings under the 150ng cDNA transfection condition, whereas the lower panel corresponds to the 250ng condition. Normalized ClC-1 current level: A531V, 1.0±0.38; A531S, 5.58±1.90; A531C, 4.67±1.76; A531G, 2.99±1.40; A531T, 2.16±0.94; A531M, 0.70±0.17; A531I, 0.64±0.32; A531E, 0.47±0.22; A531F, 0.46±0.27; A531F, 0.46±0.27; A531Y, 0.43±0.24; A531R, 0.43±0.20; A531K, 0.44±0.16; A531D, 0.40±0.13; A531L, 0.21±0.13; untransfected, 0.00 ± 0.03. Asterisks denote significant difference from the vector control (*, $P < 0.05$; n = 50-110, 22-133).

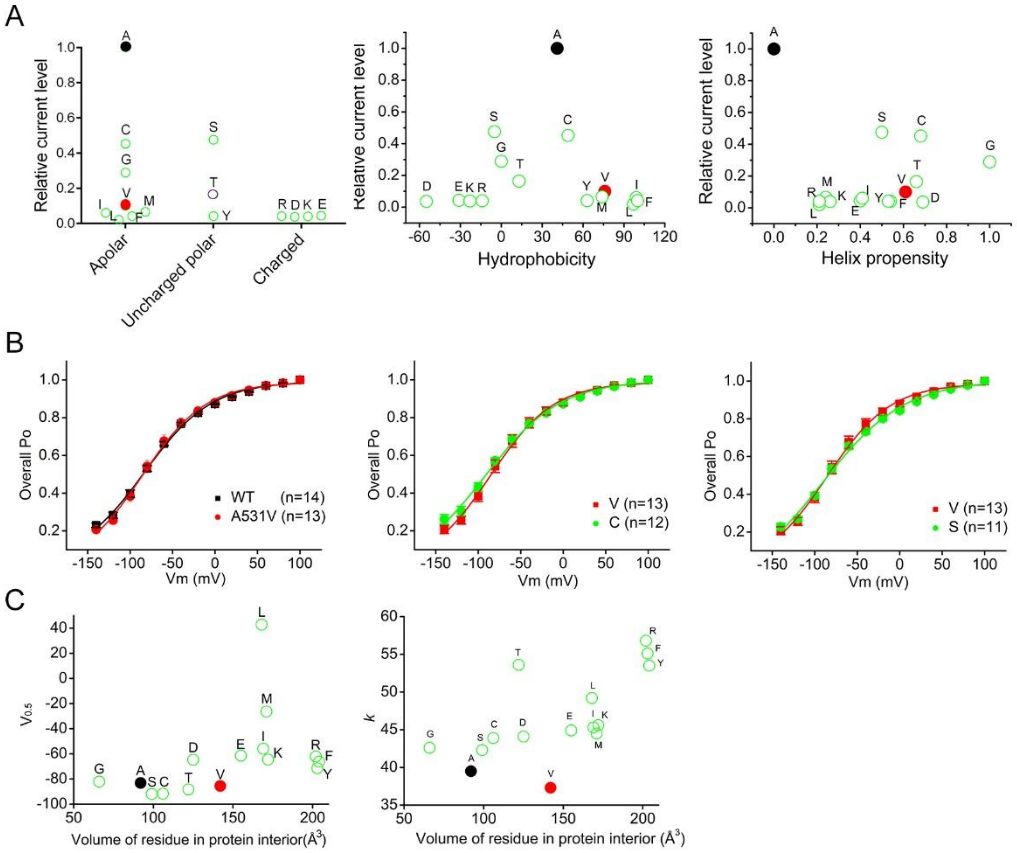


***Supplementary Figure S2. Electrophysiological analysis demonstrated that the A531 substitutions exhibited no significant difference in gating properties, and critically, no correlation was observed between the recorded current levels and various protein properties.***

(A) ClC-1 current levels of A531 mutants normalized to WT and plotted against (*left*) amino acid chemical structures (apolar, uncharged polar, charged), (*middle*) hydrophobicity, normalized to glycine (hydrophobicity index = 0), and (*right*) helix propensity, with alanine defined as 1.0 as the strongest helix-forming residue. Helix propensity values of other amino acids were derived from simulation. (B) Steady-state activation (Po–V) curves derived from isochronal tail currents at −100 mV in response to various test pulse potentials. Po–V curve comparisons of ClC-1 WT with A531V (*left*), A531V with A531C (*middle*), and A531V with A531S (*right)* revealed no significant differences. (C) The (*Left*) $V_{0.5}$ and (*Right*) $k$ values of ClC-1 A531 substitutions showed no correlation with the residue volume of the substituted amino acids. The Po–V curves generated from electrophysiological data were fitted using the Boltzmann equation, $Po(V) = P_{min} + (1- P_{min})/\{1+\exp[(V_{0.5}-V)/k]\}$, where $V_{0.5}$ represents the half-activation potential and $k$ denotes the slope factor.

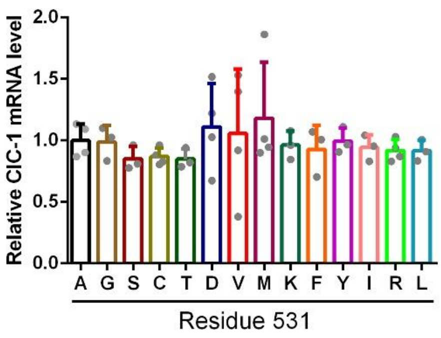


***Supplementary Figure S3. No significant differences in ClC-1 mRNA expression levels were detected among the A531 substitutions.***

mRNA levels of ClC-1 WT and A531 mutants were quantified by RT-qPCR and normalized to the corresponding ClC-1 WT control. No significant differences were detected (n = 3-4). Normalized ClC-1 mRNA level: WT, 1.0±0.13; A531G, 0.98±0.13; A531S, 0.85±0.09; A531C, 0.86±0.07; A531T, 0.84±0.08; A531D, 1.11±0.35; A531V, 1.05±0.52; A531M, 1.18±0.45; A531K, 0.96±0.11; A531F, 0.92±0.19; A531Y, 0.99±0.10; A531I, 0.94±0.10; A531R, 0.91±0.08; A531L, 0.91±0.08.

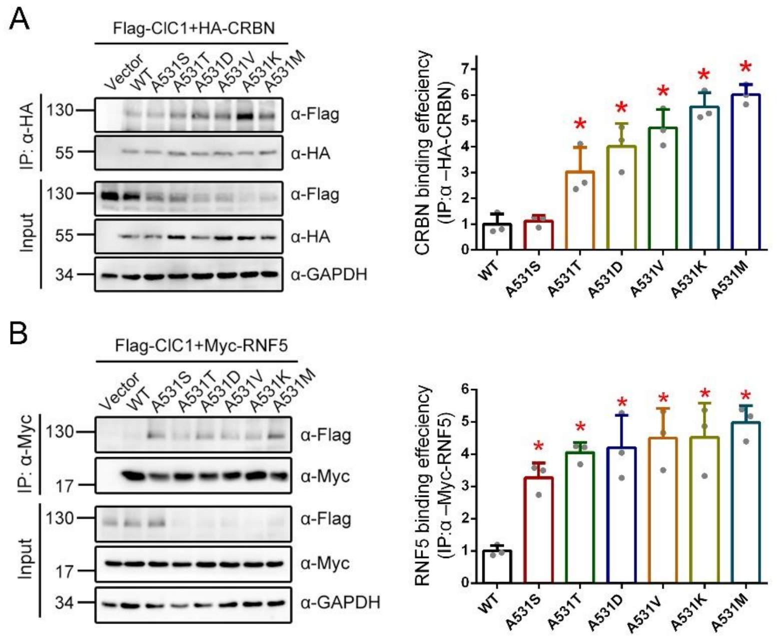


***Supplementary Figure S4. A531 substitutions strengthen the binding affinity between ClC-1 and the substrate receptor CRBN.***

Co-immunoprecipitation of ClC-1 A531 mutations and (A) HA-CRBN or (B) Myc-RNF5. Co-expression with the HA or Myc vector was used as the control. (*left panel*) Representative immunoblots of binding efficiency assay. HEK293T cell lysates were immunoprecipitated (IP) with anti-HA or anti-Myc, followed by immunoblotting with anti-Flag, anti-HA or anti-Myc. Corresponding expression levels of ClC-1 and HA-CRBN in the lysates are shown in the "input" lane. Hereafter, input represents~10%of the total protein used for immunoprecipitation. (*right panel*) Quantitative analysis of ClC-1 binding efficiency for CRBN or RNF5. Binding efficiency was calculated as the ratio of input-normalized IP ClC-1 to input-normalized IP CRBN or RNF5. These ratios were subsequently normalized against the WT control to facilitate comparison across groups. Normalized ClC-1-CRBN binding efficiency: WT, 1.0±0.39; A531S, 1.1±0.22; A531T, 3.03±0.90; A531D, 4.1±0.89; A531V, 4.73±0.72; A531K, 5.54±0.55; A531M, 6.02±0.38 (*, $P$ < 0.05; n = 3). Normalized ClC-1-RNF5 binding efficiency: WT, 1.0±0.16; A531S, 3.26±0.45; A531T, 4.04±0.31; A531D, 4.19±1.03; A531V, 4.49±0.92;

A531K, 4.51±1.06; A531M, 4.98±0.52 (*, $P < 0.05$; n = 3)

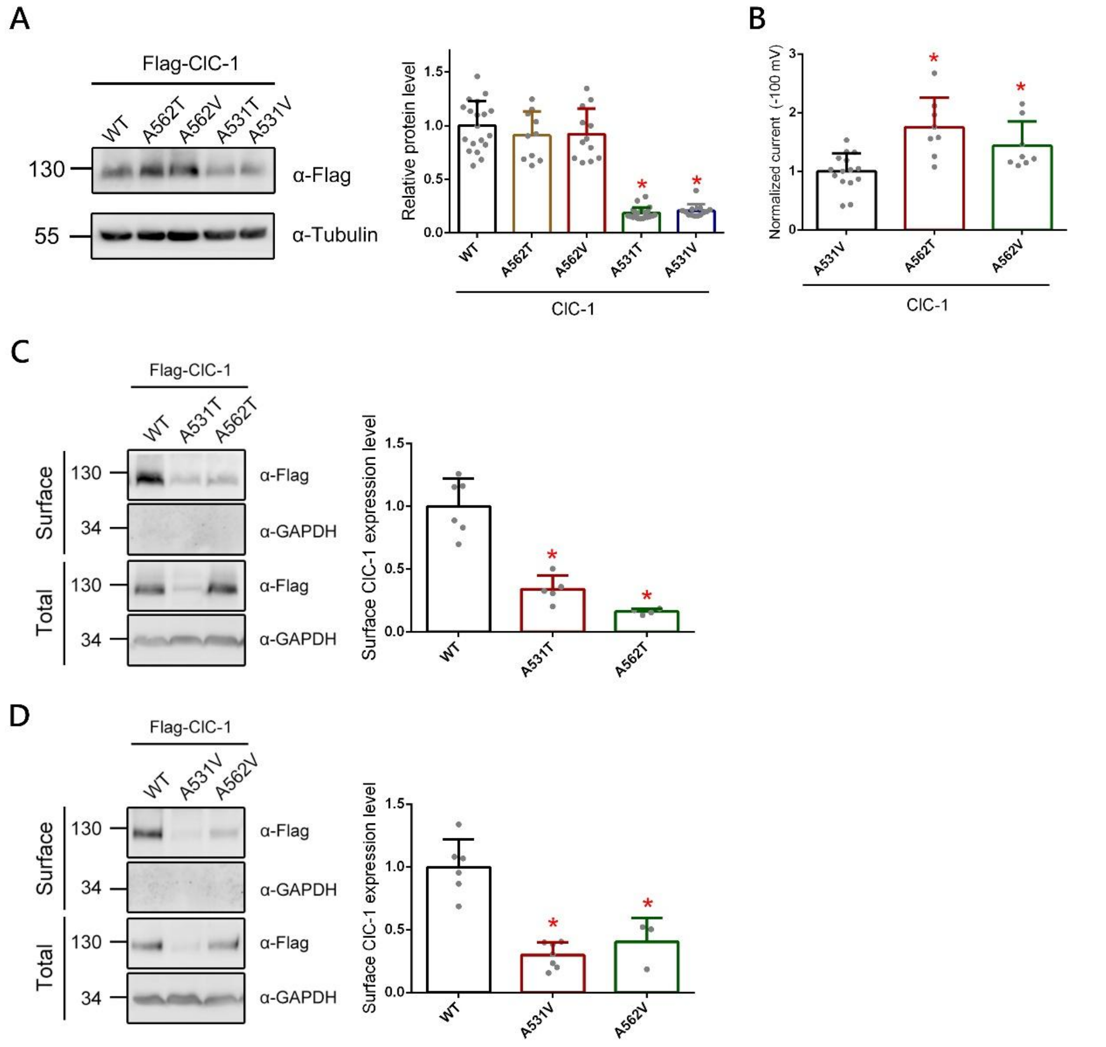


***Supplementary Figure S5. The A562 mutation showed no significant effect on ClC-1 protein expression***

(A) (*Left*) Immunoblot analysis and (*Right*) quantification of ClC-1 wild type and mutants in which alanine at residue 531 was substituted with threonine or valine, as well as the nearby residue A562 (*, $P < 0.05$; n = 10-21). (B) Quantification of ClC-1 current recordings using whole-cell patch clamp. Steady-state currents at –100 mV were normalized to the corresponding A531V control. Normalized ClC-1 current level: A531V, 1.0±0.31; A562T, 1.74±0.50; A562V, 1.43±0.41 (*, $P < 0.05$; n = 8-15). Representative immunoblots and quantitative analysis of biotinylated HEK293T cell lysates expressing (C) ClC-1-A531T and A562T, (D) ClC-1-A531V and A562V. Total protein was analyzed directly, whereas surface protein was enriched by streptavidin pull-down prior to immunoblotting. GAPDH was included as a loading control. Surface protein intensities were first standardized to the corresponding total GAPDH signals and subsequently normalized to the WT control. Normalized surface ClC-1 protein level: (C) WT, 1.0±0.22; A531T, 0.34±0.10; A562T, 0.16±0.02; (D) WT, 1.0±0.22; A531V, 0.29±0.10; A562V, 0.40±0.18 (*, $P < 0.05$; n = 3-6).

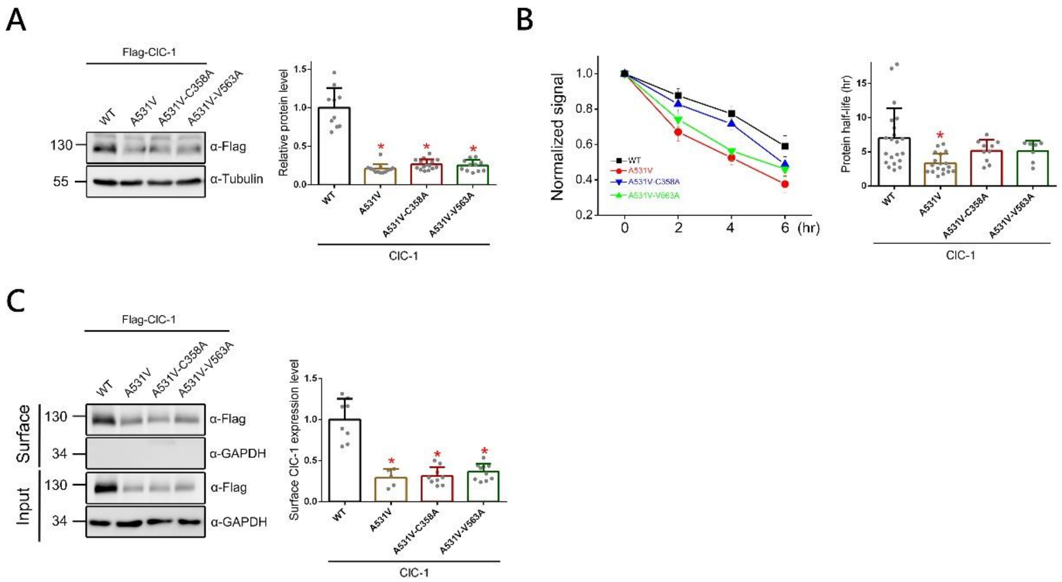


***Supplementary Figure S6. The spatial rearrangement adjacent to the ClC-1 A531V mutation did not restore protein stability or surface expression***

(A) Representative immunoblots and quantification of ClC-1 A531V mutant compare to double mutants. Normalized ClC-1 protein level: WT, 1.0±0.25; A531V, 0.20±0.06; A531V-C358A, 0.26±0.06; A531V-V563A, 0.25±0.07 (*, $P$ < 0.05; n = 10-15). (B) (*Left*) Linear plot showing the relative protein levels of ClC-1 protein following various durations of cycloheximide (CHX) treatment. Protein signals were quantified as the ratio of ClC-1 to the corresponding α-Tubulin signals, and subsequently normalized to the untreated control (0 h). Transfected HEK293T cells were subjected to 100 μg/ml CHX treatment for indicated durations (0-6h) (*Right*) Quantitative analysis of ClC-1 protein half-life in the presence of A531V or other double mutations. Normalized ClC-1 protein half-life: WT, 6.97±4.38; A531V, 3.30±1.44; A531V-C358A, 5.16±1.59; A531V-V563A, 5.10±1.51 (*, $P$ < 0.05; n = 7-21). (C) Surface biotinylation analysis showing that ClC-1 double mutants failed to restore cell-surface protein levels compared with the ClC-1 A531V mutant. (*Left*) Representative immunoblots of biotinylated HEK293T cell lysates. Total protein was analyzed directly, whereas surface protein was enriched by streptavidin pull-down prior to immunoblotting. GAPDH was included as a loading control. (*Right*) Quantification of surface protein expression. Surface protein intensities were first standardized to the corresponding total GAPDH signals and subsequently normalized to the WT control. Normalized surface ClC-1 protein level: WT, 1.0±0.25; A531V, 0.29±0.10; A531V-C358A, 0.31±0.10; A531V-V563A, 0.36±0.09 (*, $P < 0.05$; n = 5-10).

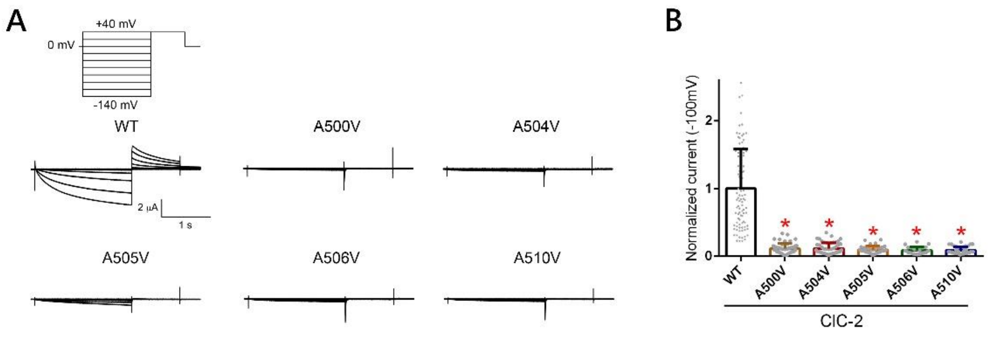


***Supplementary Figure S7. Alanine substitutions within helix O of ClC-2 resulted in reduced current levels compared with WT.***

(A) Representative current traces of ClC-2 WT, A500V, A504V, A505V, A506V and A510V. In all oocyte injection conditions hereafter, the cRNA concentration is 5 ng/µl for each construct. The external bath solution contains 3 mM KCl. From a holding potential of 0 mV, cells were subjected to 2 s voltage steps ranging from –140 to +40 mV in 20 mV increments, followed by a 1-s tail pulse to +40 mV and then returned to 0 mV (*Left panel*). After each sweep, the P/–2 leak subtraction protocol with a holding potential of 0 mV was applied to eliminate leak currents. After each sweep, the P/–8 leak subtraction protocol with a holding potential of 0 mV was applied to eliminate leak currents. (B) Quantification of ClC-2 current recordings in Xenopus oocytes. Steady-state currents at –100 mV were normalized to the corresponding WT control. Normalized ClC-2 current level: WT, 1.0±0.58; A500V, 0.11±0.07; A504V, 0.11±0.08; A505V, 0.09±0.05; A506V, 0.08±0.05; A510V, 0.08±0.05 (*, $P < 0.05$; n = 29-95).

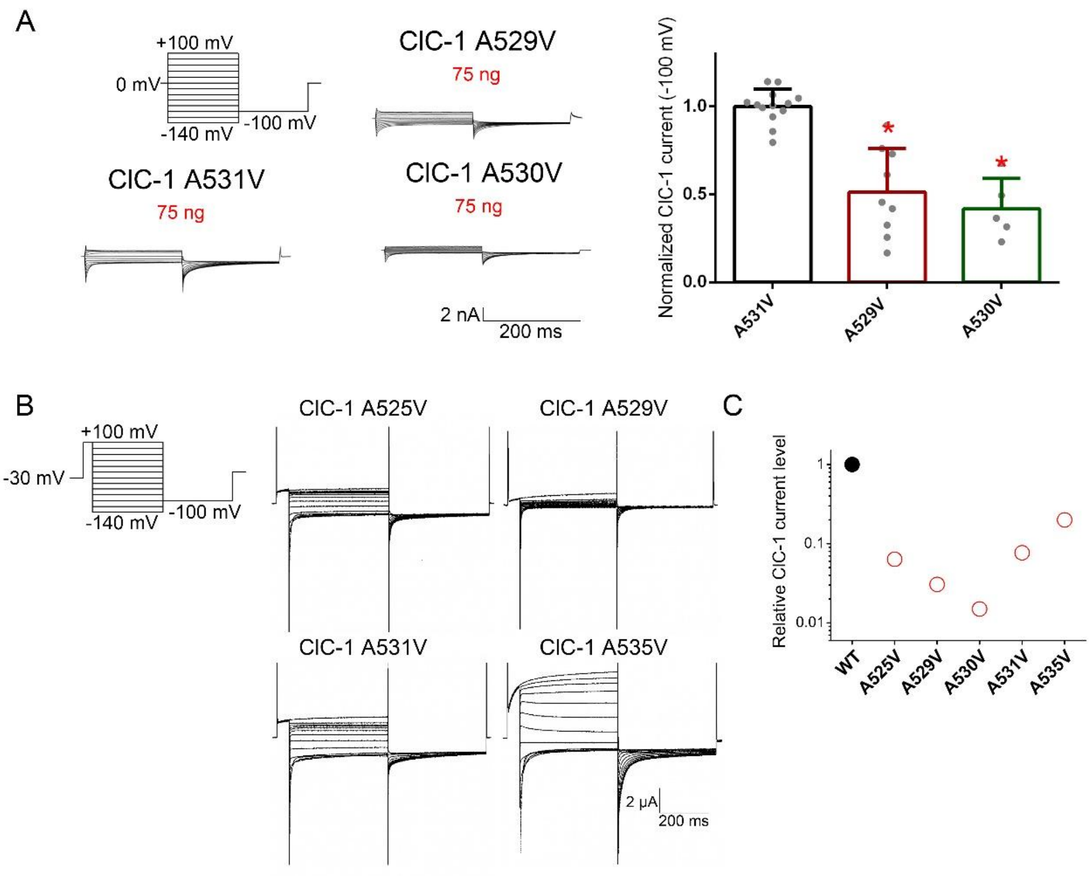


***Supplementary Figure S8. Helix O alanine substitutions resulted in a decreased ClC-1 current level***

(A) Whole-cell patch-clamp experiments performed in HEK293T cells expressing ClC-1-A531V, ClC-1-A529V and ClC-1-A530V. (*Left*) Representative $Cl^-$ current traces recorded from human ClC-1 channels. Representative $Cl^-$ current traces recorded from human ClC-1 channels. From a holding potential of 0 mV, cells were subjected to 200-ms voltage steps ranging from +100 to –140 mV in –20 mV increments, followed by a tail-voltage step to -100 mV for 200 ms, as shown in the left panel. Cells expressing ClC-1-A5331V were used as controls. (*Right*) Quantification of relative peak Cl- current amplitude at -100 mV. Data were normalized with respect to the ClC-1-A531V control. Normalized ClC-1 current amplitude: A531V, 1.0±0.09; A529V, 0.51±0.24; A530V, 0.41±0.17. Asterisks denote significant difference from the vector control (*,

*P* < 0.05; n = 5-13) (B) Representative current traces of ClC-1 WT, A525V, A529V, A531V and A535V. The cRNA concentration is 2.07 ng/μl for WT construct and 41.4 ng/μl for other mutation construct. From a holding potential of -30 mV, cells were subjected to a 50 ms prepulse at +100 mV. This was followed by 400 ms voltage steps ranging from +100 to -140 mV in -20 mV decrements. A 200 ms tail pulse to -100 mV was then applied before returning to the holding potential. (C) Quantification of ClC-1 currents recorded in HEK293T cells and Xenopus oocytes. Mutant current amplitudes were normalized to WT levels. The y-axis is presented on a semi-logarithmic scale.
